# Reliable energy-based antibody humanization and stabilization

**DOI:** 10.1101/2022.08.14.503891

**Authors:** Ariel Tennenhouse, Lev Khmelnitsky, Razi Khalaila, Noa Yeshaya, Ashish Noronha, Moshit Lindzen, Emily Makowski, Ira Zaretsky, Yael Fridmann Sirkis, Yael Galon-Wolfenson, Peter M. Tessier, Jakub Abramson, Yosef Yarden, Deborah Fass, Sarel J. Fleishman

**Author notes:** University of California San Francisco, Department of Urology, School of Medicine (Ashish Noronha).

## Abstract

Humanization is an essential step in developing animal-derived antibodies into therapeutics, and approximately one third of approved antibodies have been humanized. Conventional humanization approaches graft the complementarity-determining regions (CDRs) of the animal antibody onto several homologous human frameworks. This process, however, often drastically lowers stability and antigen binding, demanding iterative mutational fine-tuning to recover the original antibody properties. Here, we present Computational hUMan AntiBody design (CUMAb), a web-accessible method that starts from an experimental or model antibody structure, systematically grafts the animal CDRs on thousands of human frameworks, and uses Rosetta atomistic simulations to rank the designs by energy and structural integrity (http://CUMAb.weizmann.ac.il). CUMAb designs of five independent antibodies exhibit similar affinities to the parental animal antibody, and some designs show marked improvement in stability. Surprisingly, nonhomologous frameworks are often preferred to the highest-homology ones, and several CUMAb designs that use different human frameworks and differ by dozens of mutations are functionally equivalent. Thus, CUMAb presents a general and streamlined approach to optimizing antibody stability and expressibility while increasing humanness.

## Introduction

Antibodies are the largest segment of protein-based therapeutics, with over 100 antibodies in clinical use or under regulatory review^1^. Approximately one third of these antibodies were isolated from an animal source such as mouse and were humanized prior to clinical application^1^. Antibody humanization is essential to achieve three important therapeutic goals: i) recruiting the immune system through Fc effector function; ii) increasing blood circulation half-life; and iii) reducing or eliminating immunogenicity, especially for long-term (repeat dosing) treatments^2–4^.

Despite its critical role, antibody humanization has typically relied on an iterative trial-and-error process^5–7^. The first step chimerizes the animal variable fragment (Fv) with human constant domains (Fig. 1A). Next, the Fv, which comprises more than 200 amino acids and is divided into the variable light and heavy regions, is humanized. In this step, the Fv is mutated to resemble natural human antibodies, ideally without compromising antibody binding function or stability. This step is complicated by the fact that the Fv contains the so-called complementarity-determining regions (CDRs), hypervariable segments which are responsible for antigen recognition and are therefore highly sensitive to mutation (Fig. 1A). Thus, the mainstream humanization strategy grafts the parental CDRs from the animal antibody onto a human framework, typically leading to an Fv with >80% sequence identity to the human germline (compared to 50-70% identity for a mouse Fv; Supplementary Table 1). The frameworks are conventionally selected according to sequence^5^ or structural homology to the animal antibody^8^. Other methods humanize predicted immunogenic segments in the parental framework^9^ instead of the entire framework, graft residues from human frameworks onto the animal antibody in solvent-exposed positions^10^, or use structure-based calculations to revert humanizing mutations that may destabilize the Fv^11^. A common pitfall is that humanization typically leads to substantial, sometimes orders-of-magnitude, decreases in antibody expression levels, stability, and affinity^5, 12–14^. These undesirable outcomes reduce efficacy, increase production costs, and complicate drug delivery and formulation^15^. Therefore, as a rule, a third step in the humanization process is often necessary, which invovles “back mutating” positions in the humanized antibody to their animal identities, requiring lengthy design and experimental iterations^6, 13^.

**Figure 1:**
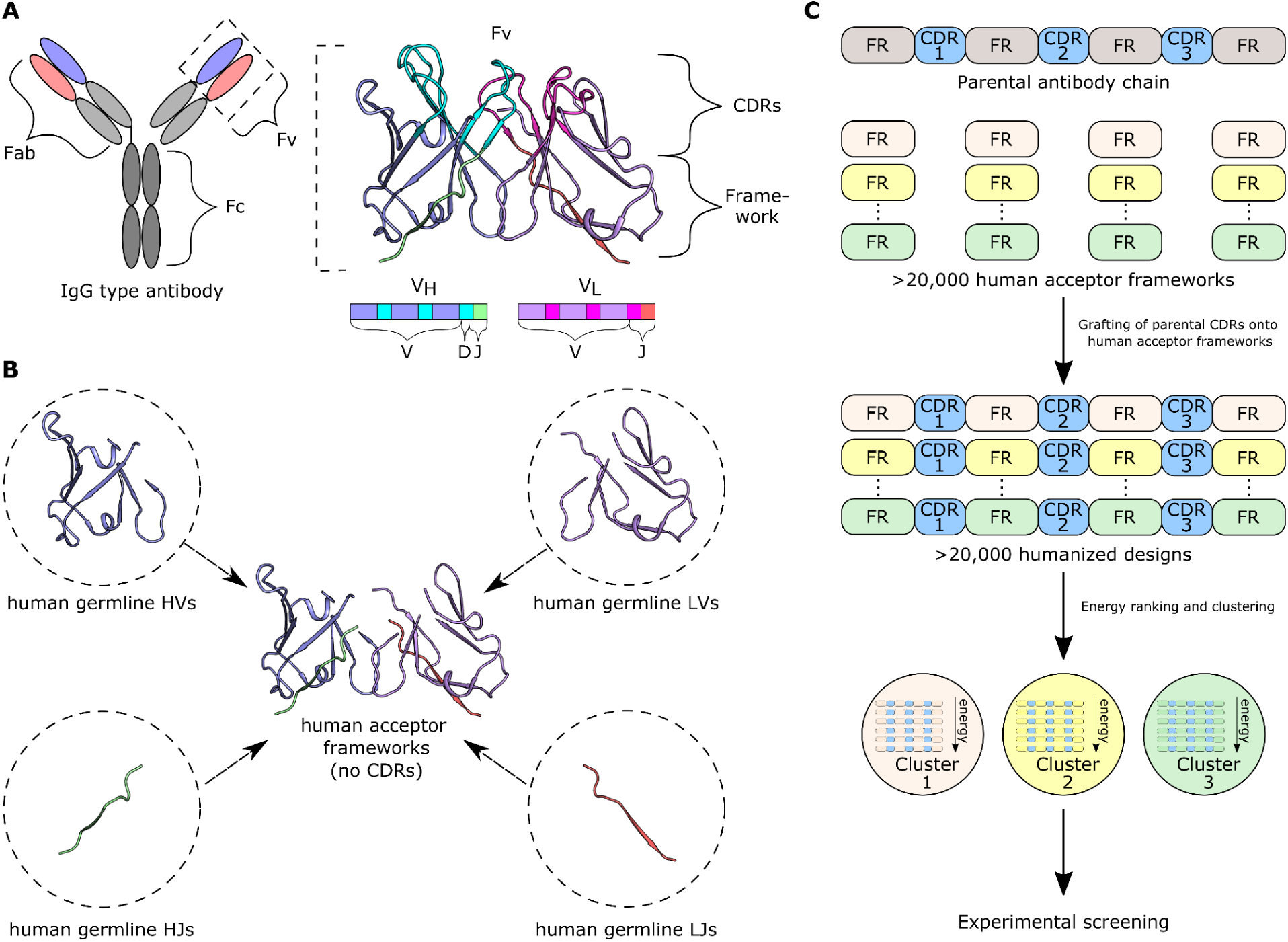
Key steps in energy-based antibody humanization using CUMAb. (**A**) Schematic of antibody domains. (**B**) CUMAb begins by combining human germline heavy and light V and J genes to generate more than 20,000 human frameworks per antibody. (**C**) CDRs from the parental antibody are grafted onto each human acceptor framework. The humanized designs are then modeled in Rosetta and clustered, yielding low-energy cluster representatives for experimental screening.

The loss in affinity and stability often caused by humanization stems from structure and energy incompatibilities between the animal CDRs and human framework^16, 17^. Previous structural analyses identified an important region in the framework, namely the vernier zone, which underlies the CDRs. This zone comprises approximately 30 positions that vary even among homologous frameworks and impact CDR structural integrity and stability^16, 17^. Thus, most back mutations are introduced to reconstitute some of the vernier-zone positions seen in the animal antibody^5^. This process can regain the parental antibody affinity and stability, though at the cost of lower humanness and lengthy iterations.

Here, we test whether energy-based ranking is more effective than homology in designing humanized antibodies. Starting from experimentally determined or computationally modeled structures, we use automated Rosetta all-atom calculations to design humanized variants and rank them solely based on their energies. Crucially, automation allows us to systematically expand the options of unique humanized designs from a few dozen per antibody using conventional humanization methods to more than 20,000. We validate the method by demonstrating that some of the top-ranked designs for five unrelated antibodies exhibit stability and binding properties on par and sometimes greater than the parental animal antibodies without requiring back mutation or iterative design/experiment cycles. Remarkably, in some cases, the experimentally best-performing humanized design derives from human framework genes that do not exhibit high homology to the animal antibody, suggesting that energy-based humanization may provide a different and more effective solution than conventional homology-based humanization. The algorithm, which we call CUMAb for Computational hUMan AntiBody design, is general, computationally efficient, and enabled for academic use through http://CUMAb.weizmann.ac.il.

## Results

### Systematic energy-based ranking of humanized antibodies

We describe a general computational workflow for modeling and energy ranking humanized Fv designs. Starting from an experimental or computational structure of the animal Fv, we replace all the framework regions with compatible human ones, leaving only the parental CDRs. The framework is encoded in two gene segments, V and J, on both the light and heavy chains (Fig. 1A)^18^. Recombining all human V and J segments on both light and heavy chains yields tens of thousands of unique frameworks (63,180 that comprise kappa light chains and 48,600 that comprise lambda light chains)^19^. Our working hypothesis is that energy-based ranking of a diverse set of human frameworks may yield humanized designs that are more stable and functional than those selected from conventional, homology-based humanization.

We defined the CDRs (Supplementary Table 2) based on visual inspection of representative Fv structures and guided by previous analyses^20, 21^, and we used this definition for all targets reported here without additional intervention. The first step in CUMAb combines all human V and J gene sequences obtained from the ImMunoGeneTics (IMGT) database^19^ (Fig. 1B) to generate acceptor framework sequences. D genes are fully encompassed by CDR H3, which is held fixed in all calculations. Kappa or lambda light chain genes are used based on the light-chain class of the animal antibody. We exclude human genes that contain Asn-Gly or Asn-X-Ser/Thr (where X is not Pro) sequence motifs because they may lead to undesirable post-translational modifications, including N-linked glycosylation^5^. Additionally, we exclude any V gene that exhibits more than two cysteines outside the CDRs to reduce the chances of aggregation or disulfide shuffling. These restrictions retain a large fraction of possible combinations of human genes, resulting in >20,000 unique human acceptor frameworks per antibody. We next concatenate the mouse CDR sequences with the human framework regions to generate humanized sequences (Fig. 1C).

Using the parental antibody Fv structure as a template, each humanized sequence is modeled using Rosetta all-atom calculations^22^. The resulting model structure is relaxed through cycles of Rosetta combinatorial sidechain packing and constrained whole-protein minimization in the entire Fv. If an experimentally determined structure of the antigen-antibody complex is available, the antigen-binding residues on the antibody are held fixed in all design calculations. Each model is ranked using the Rosetta all-atom energy function 2015 (ref2015), which is dominated by van der Waals interactions, hydrogen bonding, electrostatics, and implicit solvation^23^. Designs in which any of the CDR backbone conformations deviate by more than 0.5 Å from the parental conformation are eliminated to ensure the structural integrity of the CDRs in their humanized context. To select a diverse set of sequences for experimental testing, we cluster the top-ranked designs according to V-gene subgroups^19^, which are defined according to sequence homology (7, 6, and 10 subgroups for heavy V, kappa, and lambda V genes, respectively). Clustering yields a shortlist of diverse, low-energy models for experimental testing. Of note, unlike conventional CDR-grafting methods, our approach is agnostic to homology between the animal and human framework. Moreover, CUMAb is scalable, as designing and ranking 20,000 different humanized constructs can be performed on our computational cluster within a few hours. RosettaScripts and command lines for running CUMAb are available as supplemental files (see code availability section).

### One-shot humanization matches parental antibody affinity on multiple frameworks

As the first target for humanization, we chose a murine monoclonal antibody mAb492.1 (mɑQSOX1) that inhibits the enzymatic activity of human Quiescin Sulfhydryl Oxidase 1 (QSOX1)^24^. This antibody is being developed as a potential cancer therapeutic due to its ability to block the contribution of QSOX1 to extracellular matrix support of tumor growth and metastasis^25^. This antibody presents a stringent test for humanization because chimerizing its mouse Fv with a human IgG1 constant region led to a complete loss of expressibility in HEK293 cells^14^. We previously addressed this challenge with the AbLIFT method, which uses atomistic design calculations to improve the molecular interactions between the Fv light and heavy domains^14^. One of the AbLIFT designs, AbLIFT18, restored expression in HEK293 cells of the chimeric antibody composed of human IgG1 constant regions and murine Fv, and this chimeric construct inhibited QSOX1 activity. This design, however, was an incompletely humanized antibody (66% and 57% V gene sequence identity to the nearest human germline in the light and heavy chains, respectively) and was not suitable for therapeutic applications in humans.

To simulate a realistic humanization scenario, CUMAb was performed directly on the parental mouse antibody mɑQSOX1. We produced expression vectors encoding the five top-ranked CUMAb designs formatted as separate light and heavy chains and experimentally tested all 15 unique pairs of these chains (five light and three heavy chains). Remarkably, 12 pairs showed comparable expression levels to AbLIFT18 by dot-blot analysis (Supplementary Fig. 1A), while no expression was detected for the chimeric construct comprising the mouse Fv and human constant domains^14^. Furthermore, electrophoretic-mobility analysis after purification indicated that seven designs showed comparable expression levels to AbLIFT18 without obvious misfolding or aggregation (Fig. 2A). The expressible designs were purified and screened for QSOX1 inhibition, and many were functional (Supplementary Fig. 1B). Two highly expressing, functional designs (hɑQSOX1.1, hɑQSOX1.2) exhibited similar inhibition constants to that of the parental antibody (Fig. 2B, Supplementary Fig. 1C). We noted that CUMAb ranked highly multiple genes in the VH3 subgroup from which hɑQSOX1.1 derives, each pairing with a different kappa light chain gene. We therefore tested two additional constructs (hɑQSOX1.3 and hɑQSOX1.4) and found hat they also exhibited similar inhibition constants to that of the parental antibody (Fig. 2B). All four designs have similar sequence identity to the human V gene germline in both the light (78-81%) and heavy (80-86%) chains. These V gene sequence identities are significantly higher than those for the mouse antibody and AbLIFT18, which both have 66% (light chain) and 57% (heavy chain) sequence identities (Fig. 2C). Additionally, the V gene sequence identities in the designs are similar to those of FDA-approved humanized antibodies, which have a mean sequence identity of 84% in the light chain and 81% in the heavy chain (Supplementary Table 1). Furthermore, the Hu-mAb^26^ and BioPhi^27^ web servers categorized all the designs as humanized (Supplementary Data Sets 1, 2). Thus, the four designs recapitulated the parental mouse expression levels and activity using different human Fv frameworks without requiring back mutations.

**Figure 2:**
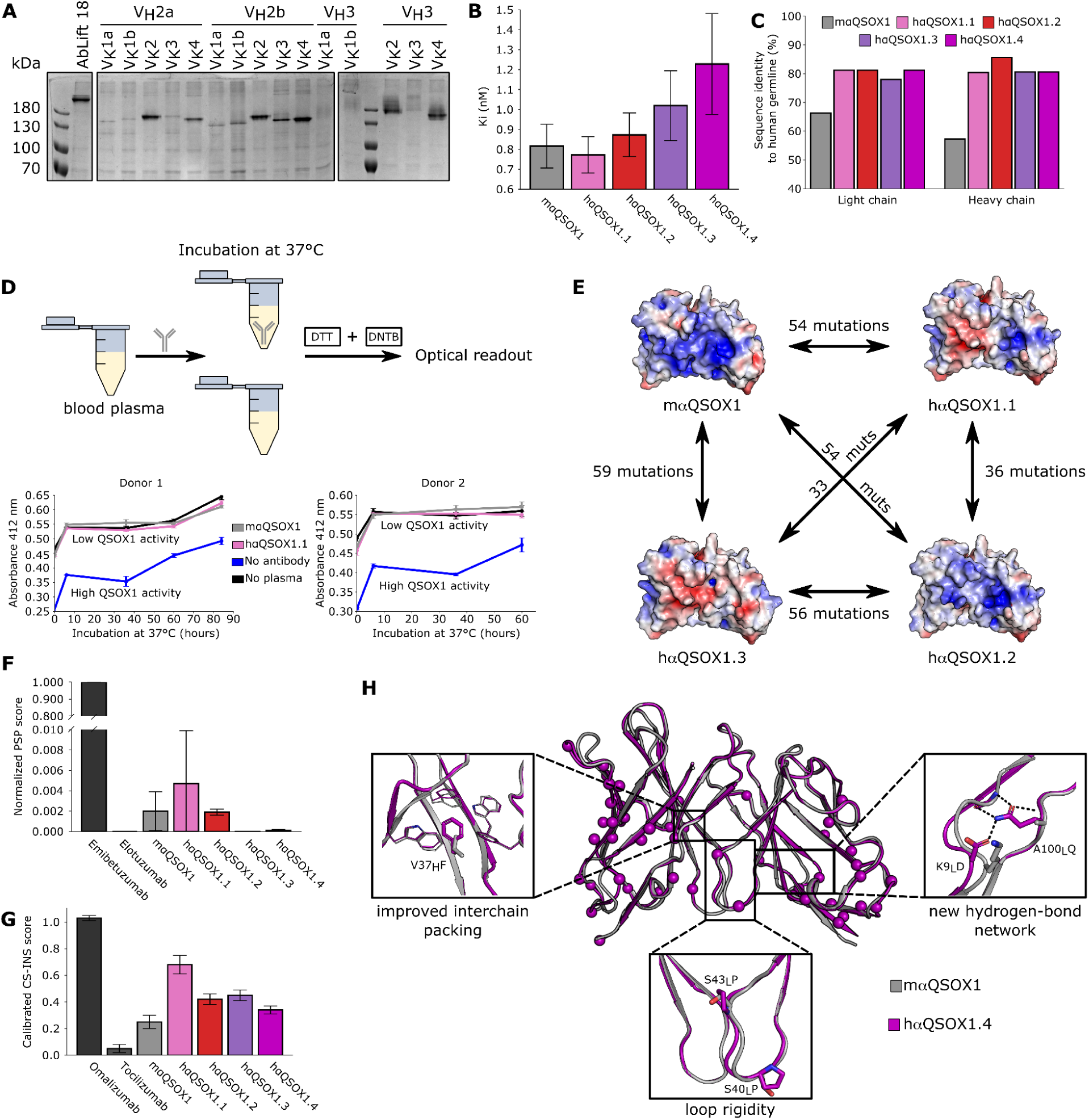
Humanization of an anti-QSOX1 antibody. (**A**) Electrophoretic analysis of 15 CUMAb designs (all combinations of the five light chains and three heavy chains from the top five clustered designs) as well as the most successful AbLIFT design (AbLift18) as a positive control^14^. CUMAb designs are labeled according to heavy chain and then by light chain, with the number referring to the IMGT subgroup from which the designed chain derives. If there are two chains from the same subgroup, they are arbitrarily labeled “a” or “b”. Amino acid sequences of each antibody are found in Supplementary Data Set 3. (**B**) K_i_ values obtained from measurements of relative QSOX1 activity at different antibody concentrations (Fits shown in Supplementary Fig. 1C). Bars shown represent the mean values, and error bars represent standard deviations of three independent repeats. (**C**) Sequence identity to the nearest human germline as calculated by IgBLAST for four CUMAb designs as well as the parental mouse antibody. (**D**) Top: Schematic of the experimental workflow. Blood plasma from human donors is incubated with an antibody at 37°C. At various time points, samples are incubated with dithiothreitol (DTT). After an incubation period to enable any uninhibited QSOX1 to oxidize DTT, 5,5′-dithiobis-(2-nitrobenzoic acid) (DTNB) was added to quantify remaining thiol groups. The colorimetric output of this assay is detected at 412 nm. Bottom: Absorbance at 412 nm at different timepoints of human blood plasma incubation with hɑQSOX1.1 (pink), mɑQSOX1 (gray), and PBS (blue) for two human blood donors, as well as PBS incubated with mɑQSOX1 (“no plasma”, black). Both hɑQSOX1.1 and mɑQSOX1 completely inhibit human QSOX1 in blood plasma at all time points. (**E**) Surface electrostatic charge of four CUMAb designs and the parental mouse antibody, showing that each design has a different electrostatic surface and that all four differ from the mouse antibody. Surface electrostatics were generated from CUMAb models using the PyMOL APBS Electrostatics plugin. (**F**) PolySpecificity Particle (PSP) measurements of nonspecific binding for mɑQSOX1 and the four CUMAb designs. Bars represent the mean values, and the error bars show the standard deviations for three independent repeats. The PSP score is normalized to a scale from 1 to 0 with 1 corresponding to high levels of nonspecific binding (emibetuzumab) and 0 corresponding to low levels of non-specific binding (elotuzumab), as described previously^29^. (**G**) Charge-stabilized self-interaction nanoparticle spectroscopy (CS-SINS) measurements of self-association for mɑQSOX1 and the four CUMAb designs in a common formulation condition (pH 6, 10 mM histidine). Bars represent the mean values, and the error bars show the standard deviations of three independent repeats. Additionally, CS-SINS results are shown for an antibody with high self-association (omalizumab) and one with low self association (tocilizumab), as described previously^28^. (**H**) Alignment of the crystal structure of the hɑQSOX1.4 Fv (magenta; PDB entry 8AON) to the crystal structure of the parental mouse antibody (gray; PDB entry 4IJ3), showing that the structures are nearly identical despite 51 mutations between them. Mutations are shown in magenta spheres. Fv structures were extracted from co-crystal structures of the antibodies with the human QSOX1 oxidoreductase fragment (QSOX1trx). The complex structure is shown in Supplementary Fig. 1E.

Next, we measured the apparent melting temperatures of the four designs as well as the parental antibody with nano-differential scanning fluorimetry (nano-DSF) and found all five to be above 70 ℃ (Supplementary Fig. 1D). To probe the stability of the strongest-inhibiting antibody in a therapeutic setting, we incubated hɑQSOX1.1 and mɑQSOX1 in human plasma at 37 ℃ and measured the inhibition of native QSOX1 as a function of time. We found that both antibodies nearly completely inhibited QSOX1 for over 60 h (Fig. 2D), indicating that the humanized designs retained the function and stability properties of the parental mouse antibody. hɑQSOX1.1, hɑQSOX1.2, and hɑQSOX1.3 differ from one another in either the light or heavy chain subgroup and have 33-56 mutations relative to one another (Fig. 2E). Due to these mutations, they have strikingly different patterns of surface charge (Fig. 2E). For instance, each pair of designs exhibits 8-15 positions with different charges (Supplementary Table 3). Thus, in this case, CUMAb produces several antibodies that are stable and functionally nearly identical but have distinct surface properties. As surface properties are associated with changes in the propensity of antibodies to self-associate or form non-specific interactions^28, 29^, CUMAb may provide a path to improving properties that are critical for therapeutic development while improving humanness. To test this hypothesis, we evaluated the non-specific binding and self-association of the four CUMAb designs and the parental antibody. Encouragingly, all five antibodies displayed low non-specific binding to a standard polyspecificity reagent^29, 30^ which correlates with low risk for abnormal pharmacokinetics and fast antibody clearance *in vivo*^31, 32^ (Fig. 2F). Moreover, all of the CUMAb designs displayed lower self-association than an FDA-approved antibody drug (omalizumab) in a common therapeutic formulation condition (pH 6, 10 mM histidine)^33^(Fig. 2G). Furthermore, one of the CUMAb designs (hαQSOX1.4) displayed lower self-association than the previously reported cutoff (CS-SINS score of <0.35)^28^, indicating that it has low risk for high viscosity or opalescence when concentrated to 150 mg/mL^33^. Thus, in this case, CUMAb produced multiple solutions with equal antigen-inhibitory activities and a range of favorable developability properties.

Finally, we determined the structure of hɑQSOX1.4 bound to the oxidoreductase fragment of human QSOX1 by x-ray crystallography and found that the design and parental antibody are strikingly similar, with only a 0.7 Å Cα RMSD despite 51 mutations between the parental and humanized antibody (Fig. 2H, Supplementary Fig. 1E, Supplementary Table 4). These results verify the atomic accuracy of CUMAb and its ability to rapidly produce functionally similar and stable humanized designs.

### Successful humanization based on Fv model structures

Experimental structure determination is a lengthy and uncertain process, and obtaining an experimentally determined structure for each isolated antibody in an antibody development campaign is unrealistic. Recently, structure-prediction methods have reached the point at which the antibody Fv structure can be predicted to nearly atomic accuracy except in the solvent-exposed parts of CDR H3, where models are still insufficiently reliable^34–39^. While H3 is a critical determinant of antigen recognition, most of its solvent-exposed region does not form direct interactions with the framework region which is the target of humanization. Therefore, we hypothesized that despite the demand for stringent accuracy by atomistic design calculations, Fv model structures might be sufficiently accurate for the purposes of humanization design. Given the recent revolution in deep-learning-based protein structure prediction methods, we chose to use AlphaFold-Multimer to predict the Fv structure of our target antibody^40, 41^. Starting from the top-ranked AlphaFold model, all design calculations are identical to the CUMAb workflow above, except that we do not eliminate humanized models in which the backbone deviates from the starting model structure, thus allowing for flexibility or modeling inaccuracies.

To test the ability of CUMAb to humanize antibodies without recourse to experimentally determined structures, we selected an antibody that was raised in mice immunized against the human AXL receptor tyrosine kinase (mɑAXL). This antibody has a framework mutation from the mouse germline that is predicted to interact with CDRH3 (Fig. 3A) and may impact its stability. Since this mutation is in the framework, CUMAb would revert it to the wild type identity in the human germline, potentially destabilizing CDRH3. To account for the uncertainty regarding this mutation, we included five designs with this mutation in addition to the five CUMAb designs. The ten designs were screened for expression, and four designs exhibited high expression levels (three designs with the additional mutation expressed but at lower levels, Fig. 3B). All ten designs were assayed for binding to AXL by immunoprecipitation, and one design showed much stronger binding than the others. This design (hɑAXL) had 40 mutations from the parental antibody and 84% (light chain) and 89% (heavy chain) V gene sequence identity to the closest human germline, compared to 62% (light chain) and 79% (heavy chain) in the mouse antibody (Fig. 3C, D). hɑAXL was also given high humanness scores by both Hu-mAb^26^ (Supplementary Data Set 1) and BioPhi^27^ (Supplementary Data Set 2). The design and the parental antibody exhibited similar affinity to AXL according to surface plasmon resonance (3.2 and 0.95 nM, respectively; Fig. 3E), and the design exhibited 3°C higher thermal stability than the parental antibody (Fig. 3F).

**Figure 3:**
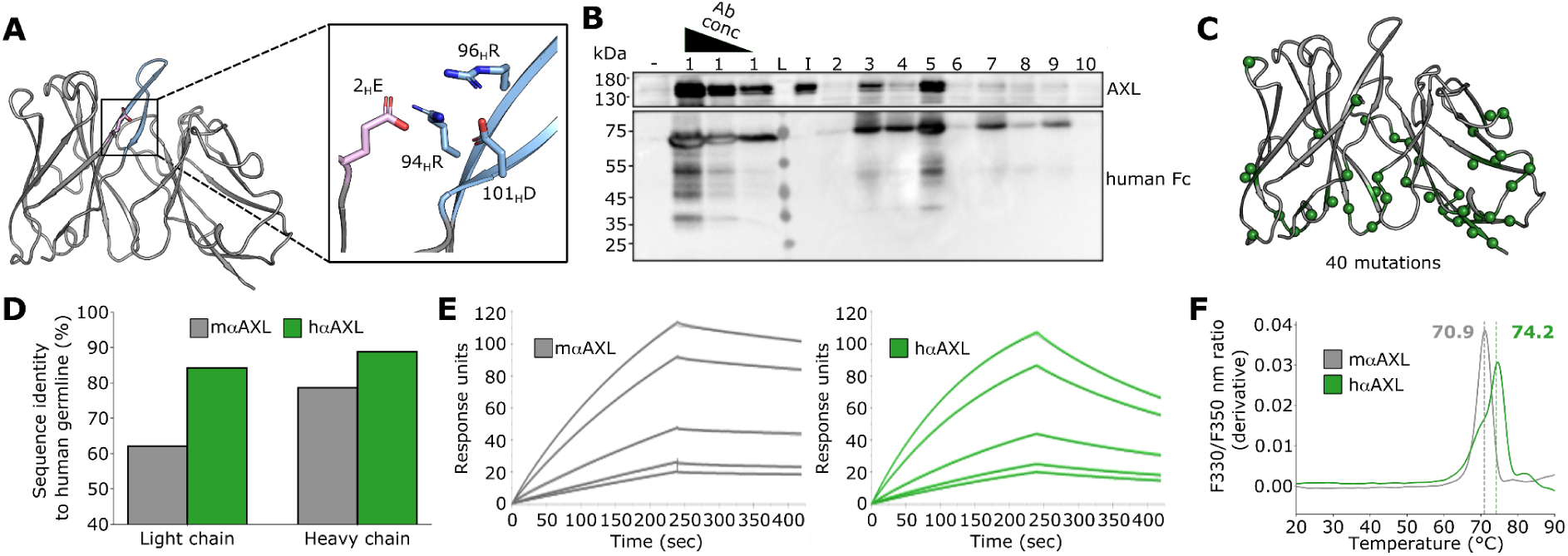
Humanization of an anti-human AXL antibody based on an AlphaFold model structure. (**A**) Unusual mutation from germline in the framework of parental mouse antibody (pink), which is predicted to interact with CDR H3 (blue). Numbering is according to Kabat numbering scheme. (**B**) Clear cell extracts were subjected to immunoprecipitation with media containing humanized AXL antibody (or control, only media) and immunoprecipitates were immunoblotted (IB) as indicated. Marked are the 10 CUMAb designs (6-10 include unusual framework mutation from A, the protein ladder (L), and the AXL protein used as input (I). (**C**) AlphaFold model of mouse anti-AXL antibody (gray) with CUMAb mutations shown in green spheres. (**D**) Sequence identity to nearest human germline as calculated by IgBLAST for the parental mouse antibody (mɑAXL, gray) as well as the best-performing CUMAb design (hɑAXL, green). (**E**) SPR kinetic analysis of mɑAXL (gray) and hɑAXL (green) to AXL (kinetic fits shown in black). Antibodies were covalently attached to a CM5 chip, and AXL was injected at 7, 5, 2, 1, and 0.8 nM. mɑAXL exhibited k_a_ = 5.5 * 10^5^ M^-1^s^-1^, k_d_ = 5.24 * 10^-4^ s^-1^, and K_D_ = 0.95 nM. hɑAXL exhibited k_a_ = 1.2 * 10^6^ M^-1^s^-1^, k_d_ = 3.9 * 10^-3^ s^-1^, and K_D_ = 3.2 nM. (**F**) Thermal denaturation of mɑAXL (gray) and hɑAXL (green) using nano-DSF. Shown is average of two independent repeats.

To further confirm the ability to humanize antibodies based on an Fv model structure, we targeted two murine antibodies (designated here as ɑMUC16_Ab1 and ɑMUC16_Ab2) against human mucin 16 (MUC16). MUC16 is a member of the mucin family that is overexpressed in ovarian cancer^42^. CA125, a cleaved soluble antigenic fragment from the tandem repeat region of MUC16, is the best-known biomarker of epithelial ovarian cancer (EOC) and is routinely monitored in patients with EOC as a prognostic tool for cancer recurrence^43^. Thus, MUC16 is considered a potential target for EOC therapeutic intervention by immunotherapy. AlphaFold models of ɑMUC16_Ab2 showed an unrealistic strand swap between the Fv light and heavy chains (see Methods), so we instead predicted this Fv structure using AbPredict^38^. We experimentally tested only one design for antibody ɑMUC16_Ab1 (hɑMUC16_Ab1) and two for antibody ɑMUC16_Ab2 (hɑMUC16_Ab2.1 and hɑMUC16_Ab2.2) and compared their expression and MUC16 binding to a partially humanized version of the parental antibody in which the mouse Fv was fused to human IgG1 (chimeric version, designated here as cɑMUC16_Ab1 and cɑMUC16_Ab2).

We first compared the secretion levels of the chimeric antibodies to each of the CUMAb designs. hɑMUC16_Ab1 exhibited nearly sevenfold higher secretion levels than its chimeric counterpart (Fig. 4A). hɑMUC16_Ab2.1 and hɑMUC16_Ab2.2 both had comparable expression levels to cɑMUC16_Ab2, with hɑMUC16_Ab2.1 having slightly higher expression levels and hɑMUC16_Ab2.2 slightly lower (Fig. 4A). We then contrasted MUC16 binding in the chimeric and humanized antibodies. The three designs displayed nearly identical affinities to MUC16 as their chimeric counterparts (Fig. 4 B, C) and essentially identical staining of HEK cells expressing a MUC16 construct (Fig. 4D, E; Supplementary Fig. 2A, B). Finally, hɑMUC1616_Ab1 showed similar staining of OVCAR-3 cells, which are known to express MUC16^44^ as cɑMUC16_Ab1 (Fig. 4F, Supplementary Fig. 2C). hɑMUC16_Ab2.1, hɑMUC16_Ab2.2, and cɑMUC16_Ab2 did not show binding to OVCAR3 cells in this FACS experiment (Fig. 4G). We concluded that the humanized antibodies retained the binding characteristics of their parents while substantially increasing expression levels in one case.

**Figure 4:**
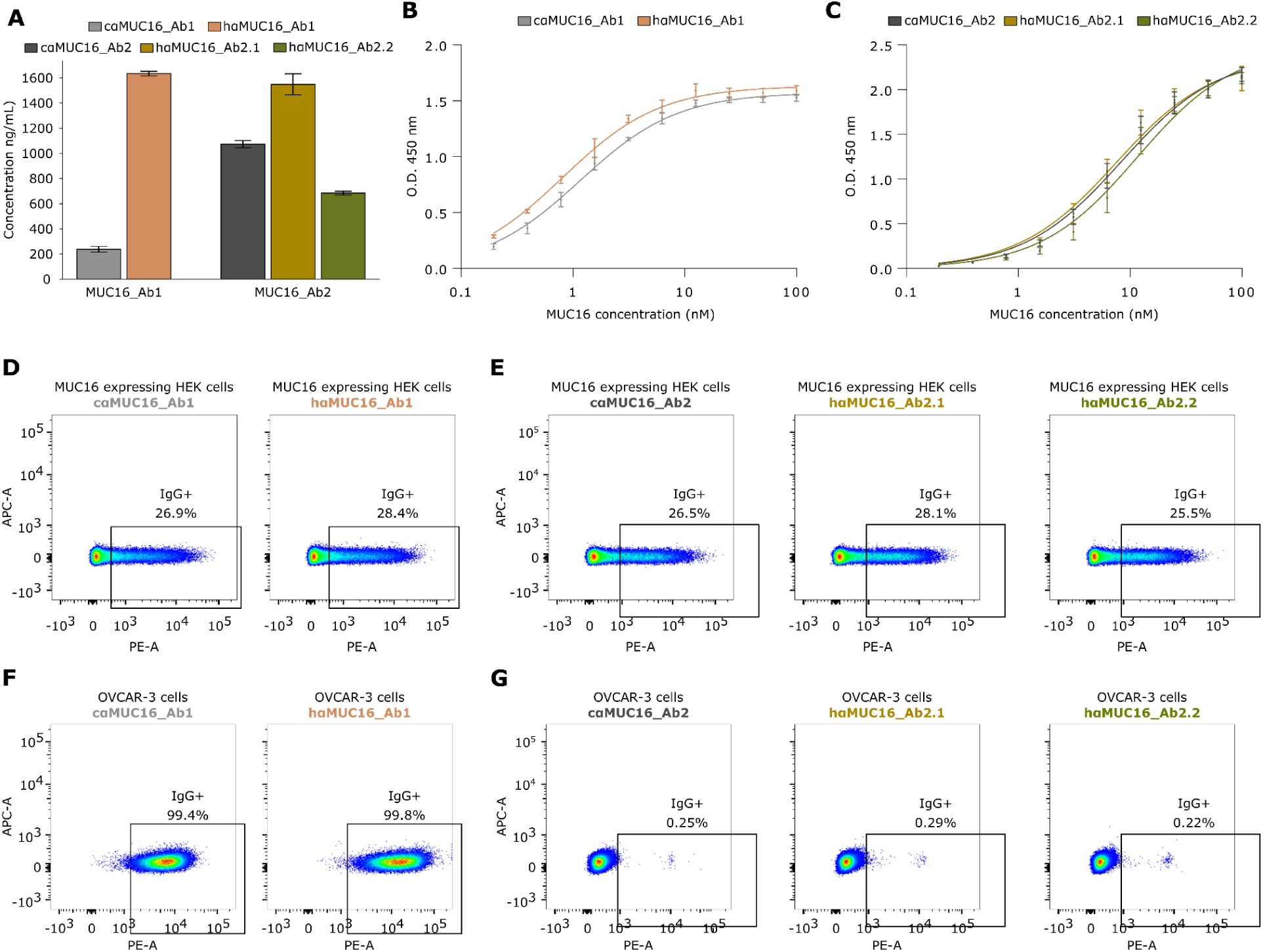
Humanization of two anti-mucin16 antibodies based on model structures. (**A**) Secretion levels of chimeric constructs and humanized designs targeting MUC16. (**B**) ELISA titration of MUC16 against constant amounts of cɑMUC16_Ab1 and hɑMUC16_Ab1. Apparent dissociation constants (K_D_): cɑMUC16_Ab1 = 1.2 ± 0.2 nM; hɑMUC16_Ab1 = 0.82 ± 0.04 nM. (**C**) ELISA titration of MUC16 against constant amount of cɑMUC16_Ab2, hɑMUC16_Ab2.1, and hɑMUC16_Ab2.2. Apparent dissociation constants (K_D_): cɑMUC16_Ab2 = 8.5 ± 1.2 nM; hɑMUC16_Ab2.1 = 7.6 ± 0.9 nM; hɑMUC16_Ab2.2 = 12.0 ± 2.5 nM. (**D-E**) HEK293 cells expressing MUC16 were labeled with cɑMUC16_Ab1 and its humanized design (D) or with cɑMUC16_Ab2 and its designs (E). (**F-G**) Acetone-fixed OVCAR-3 cells were labeled with cɑMUC16_Ab1 and its humanized design (F) or with cɑMUC16_Ab2 and its humanized designs (G). (A-C) Points or bars represent means of three independent repeats and errors bars represent standard deviations. (D-G) Relevant controls can be found in Supplementary Figure 2.

The three humanized designs have approximately 50 mutations from their respective parental antibodies and exhibit high sequence identity to the human germline (Supplementary Table 5). Strikingly, in all three cases, the successful CUMAb design derives from the human germline subgroup with the highest sequence identity to the parental antibody in only the light or heavy chain and not in both, demonstrating again that CUMAb finds solutions that conventional methods for antibody humanization would be unlikely to find (Supplementary Table 5). Additionally, hɑMUC16-2.1, and hɑMUC16-2.2 come from different light chain V gene subgroups, providing additional evidence that CUMAb is able to produce functionally identical variants coming from different frameworks.

We conclude that despite the uncertainty when using Fv model structures rather than experimentally determined ones, CUMAb can rapidly generate stable, well-expressed, high affinity humanized antibodies.

### Increased humanness through specificity-determining residue grafting

The CUMAb CDR grafting procedure typically raises the sequence identity to the human germline to 80-88% (Supplementary Table 5), in line with the levels observed for antibodies in clinical use (Supplementary Table 1). To increase humanness even further, an alternative strategy, called specificity-determining residue (SDR) grafting, was proposed^45, 46^. SDR grafting maintains only the amino acid positions that directly contact the antigen, whereas the remainder of the antibody, including CDR regions that do not interact with the antigen, is humanized. We next extend the CUMAb energy-based strategy to SDR grafting. To prevent undesirable changes to the CDR backbone conformations in SDR grafting, we use only human germline genes that exhibit CDRs that match the lengths of the CDRs in the parental antibody, with the exception of H3 (see Methods). Furthermore, because the heavy-chain J gene segment encodes a part of H3, we cluster the resulting humanized designs according to their V gene subgroups as well as their heavy-chain J segment. Thus, unlike CDR grafting above, CUMAb SDR grafting may select designs that differ from one another only in their heavy-chain J segments. We limit the application of SDR grafting to experimentally determined antigen-bound structures because this procedure demands accurate determination of antigen-binding amino acid positions, including the solvent-accessible region in H3.

We applied SDR grafting to the murine antibody D44.1 (mɑHEWL), which targets hen-egg white lysozyme. The co-crystal structure (PDB entry: 1MLC)^47^ reveals that 28 of 63 CDR positions in D44.1 interact with the antigen. Thus, in this case, SDR grafting would lead to roughly 90% (heavy chain) and 94% (light chain) V gene sequence identity to the human germline, compared to 86% (heavy chain) and 79-88% (light chain) using the CDR grafting procedure. In an initial experimental screen, we formatted six designs as single-chain variable fragments (scFvs) and tested lysozyme binding using yeast-surface display^48^ (Supplementary Fig. 3A, B). This preliminary screen indicated that five designs exhibited high expression levels and that designs 1 and 6 bound lysozyme, with design 1 markedly superior. This qualitative single-concentration experiment indicated that design 1 may bind lysozyme as strongly as the parental D44.1.

To obtain quantitative binding data, we formatted design 1 (hɑHEWL) as a human IgG1 Fab and expressed it and D44.1 (as mouse IgG1 Fab) in HEK293 cells. Design 1 has 64 mutations relative to the parental mouse antibody and substantially increases the antibody’s humanness in both the light and heavy chains (Fig. 5A, B; Supplementary Table 5). hɑHEWL is categorized as human by both Hu-mAb^26^ (Supplementary Data Set 1) and BioPhi ^27^ (Supplementary Data Set 2). Both antibodies expressed well, with the design showing noticeably higher purification yield when the Fabs were purified using an anti-His tag column (Fig. 5C). Furthermore, the melting temperature of the designed variant was 5℃ higher than that of the mouse (Fig. 5D). We measured the affinity of the mouse and humanized Fabs, and found that the humanized design has an affinity of 44 nM compared to 6.5 nM for the parental mouse antibody (Fig. 5E). Thus, in this case, the greater level of humanization enabled by SDR grafting may have come at the cost of some loss in affinity, though as with CDR grafting, expressibility and stability have increased.

**Figure 5:**
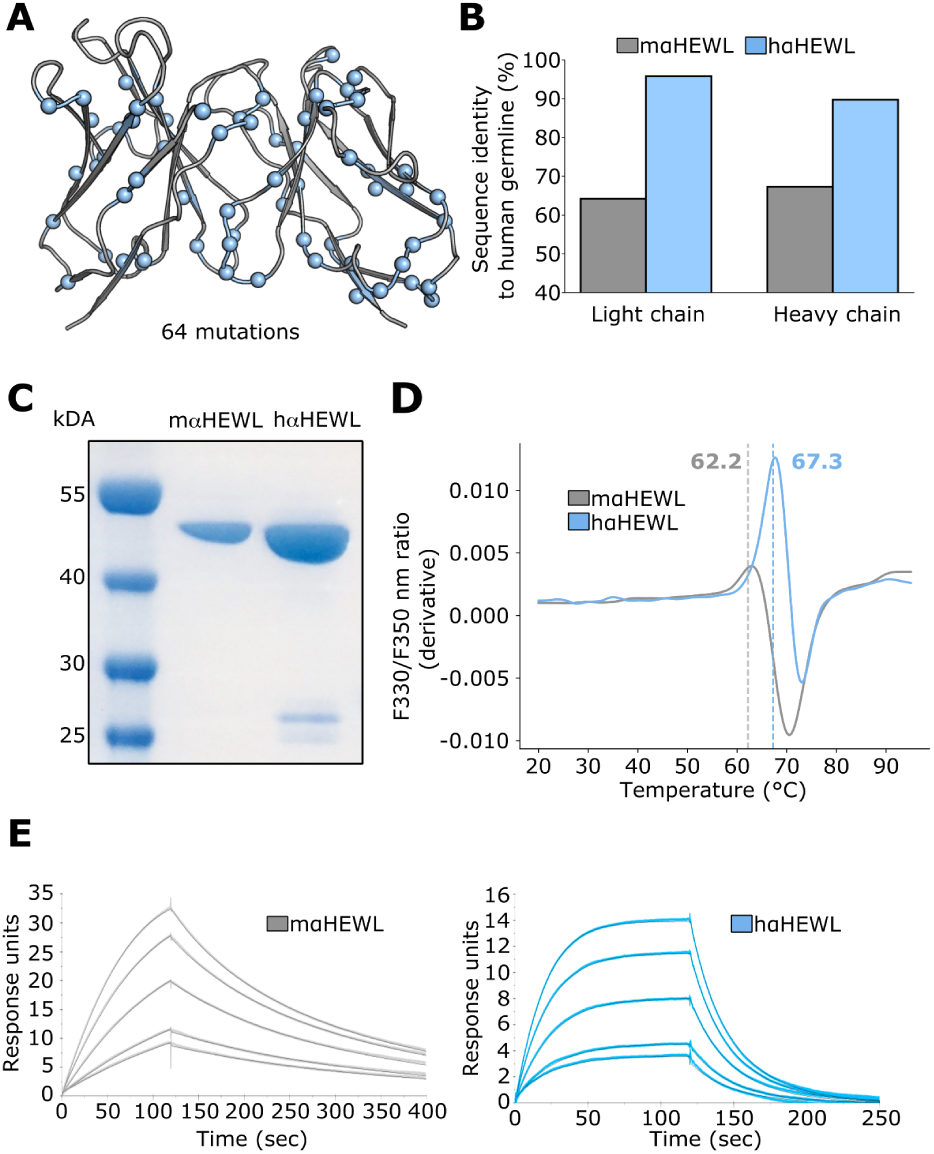
SDR-grafting of anti-hen egg-white lysozyme (HEWL) antibody. **(A)** Crystal structure of mouse anti-HEWL antibody (PDB entry: 1MLC) with CUMAb design 1 mutations shown in blue spheres (**B**) Sequence identity to nearest human germline as calculated by IgBLAST for the parental mouse antibody (gray, mɑHEWL) as well as the best-performing CUMAb design (blue, hɑHEWL). (**C**) Electrophoretic mobility analysis of mɑHEWL and hɑHEWL. Equal volumes of each protein were analyzed after purification with anti-His tag column. **(D)** Thermal denaturation of mɑHEWL (gray) and hɑHEWL (blue) using nano-DSF. Shown is average of two independent repeats. (**E**) SPR kinetic analysis of mɑHEWL (gray) and hɑHEWL (blue) to AXL (kinetic fits shown in black). Antibody Fabs were covalently attached to a CM5 chip, and HEWL was injected at 20, 15, 10, 5, and 3.75 nM. mɑHEWL exhibited k_a_ = 2.0 * 10^6^ M^-1^s^-1^, k_d_ = 1.4 * 10^-2^ s^-1^, and K_D_ = 7.4 nM. hɑAXL exhibited k_a_ = 3.9 * 10^6^ M^-1^s^-1^, k_d_ = 1.6 * 10^-1^ s^-1^, and K_D_ = 42 nM.

### Successful energy-based humanization exploits nonhomologous frameworks

Conventional antibody humanization strategies use the human germline genes that are closest to those of the sequence or structure of the animal antibody. CUMAb’s energy-based strategy ignores sequence or structure homology and focuses instead on the structural integrity and energy of the design models. Energy-based humanization vastly expands the scope of potential frameworks from a few dozen high-homology ones to tens of thousands. We analyzed the five test cases presented here to understand the role of energy versus sequence homology in successful antibody humanization.

Strikingly, in four of the five cases, the best-performing humanized designs do not derive from the best-matched human ones according to sequence identity (Supplementary Figure 4, Supplementary Table 5). In the anti-QSOX1 antibody, none of the best designs derives from the closest human V gene subgroup for either the light or heavy chain, in the anti-MUC16 antibodies, none of the three designs match their parental antibody in both the heavy and light chains, and in the anti-lysozyme antibody, only the heavy chain matches. Furthermore, the anti-lysozyme designs based on the same subgroup of human V genes were significantly inferior to the best-performing humanized design. As an additional comparison with conventional humanization strategies, we attempted to humanize the anti-QSOX1 antibody using the “consensus” approach. In this humanization approach, the framework is taken from a sequence-based consensus of V gene subgroups^49^ (in this case, IGKV1 and IGHV4). The designed antibody, however, failed to express, and we could not test its binding to QSOX1 (Supplementary Fig. 5), providing another demonstration that energy-based humanization may be superior to conventional humanization strategies.

Thus, grafting CDRs from an animal antibody onto the closest (or consensus) human germline often fails to recapitulate their stability and functional properties, as has been observed in decades of antibody engineering^12, 13^. By contrast, some of the top-ranked designs from energy-based humanization exhibit stability and binding properties indistinguishable from those of the animal antibody, and these designs typically do not derive from germline gene subgroups that are homologous to those of the animal antibody.

We next ask what are the structural underpinnings of successful design using nonhomologous frameworks. Our analysis carries significant uncertainty because the designs diverge from one another by dozens of mutations across the framework, and thus, claims on which mutations determine success or failure must be tentative. Yet, comparing the designs that use homologous frameworks with the best-performing designs shows striking structural differences that may account for the higher stability and activity of the latter. For instance, humanization of the anti-QSOX1 antibody based on the consensus framework of the VH4 subgroup, which is the highest homology subgroup according to IgBLAST, results in two mutations from the parental antibody in the framework regions. One of these mutations eliminates a buried hydrogen bond network that connects the framework and CDRs 1 and 2 (Fig. 6A) and another leads to steric overlaps with a position on CDR H1 (Fig. 6B). Additionally, all successful CUMAb designs introduced light-chain mutation Ser43Pro, which is also seen in the AbLIFT18 design and is likely to stabilize the antibody (Fig. 6C)^14^. Humanizing the anti-lysozyme antibody with the highest-homology VK3 family would result in a His34Ala mutation in the light chain, disrupting a hydrogen-bond network among the light chain CDRs (Fig. 6D). Strikingly, although hɑHEWL introduces a mutation in this same critical position (His34Asn), this mutation is predicted to retain the hydrogen-bond network. We conclude that despite the high homology within V-gene subgroups, critical long-range interactions between the framework and the CDRs are often abrogated by conventional homology-based grafting. By contrast, low-energy ranked designs often retain the very same identities seen in the parental antibody and sometimes mutate them while nevertheless conserving critical stabilizing interactions.

**Figure 6:**
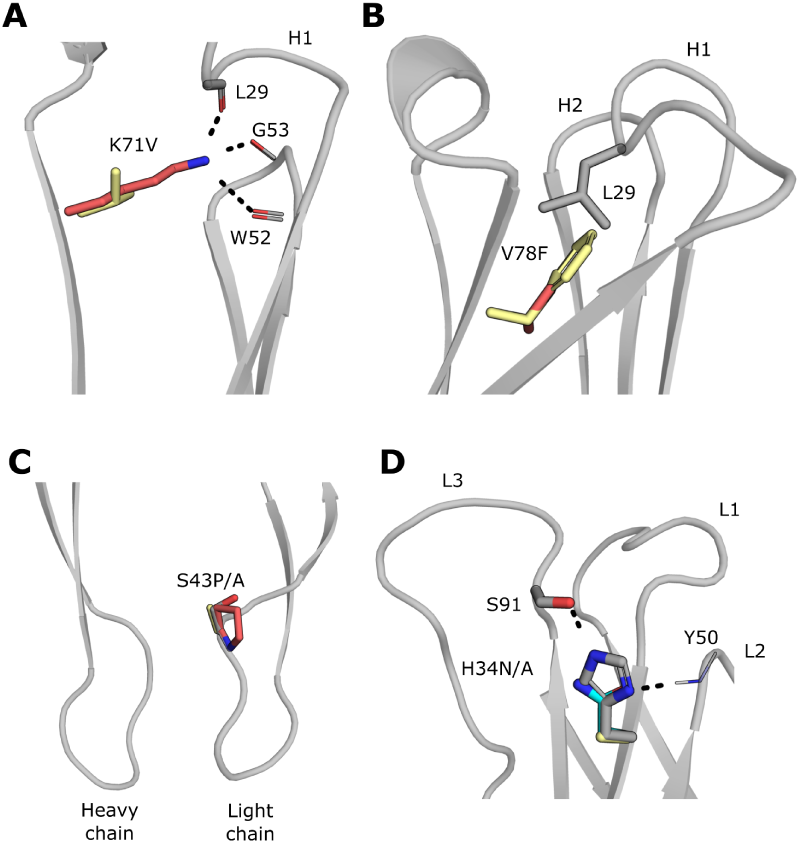
Structural determinants of successful humanization by nonhomologous frameworks. **(A-C).** Crystal structure of mouse anti-QSOX1 (gray, PDB entry: 4IJ3) aligned with CUMAb model of successful design hɑQSOX1.2 (red) and CUMAb model of unsuccessful humanization onto consensus framework IGKV1-IGHV4 (yellow). Amino acid numbering according to Kabat^20^. **(A)** Lys71 in the heavy chain framework of the mouse anti-QSOX1 antibody makes three stabilizing hydrogen bonds with backbone carbonyls of CDRs H1 and H2. This Lys is conserved in the successful design but mutated to Val in the consensus framework IGHV4, likely destabilizing the H1 and H2 loops. **(B)** Val78 in the heavy chain framework of mouse anti-QSOX1 is mutated to Phe in the consensus framework IGHV4. This likely strains the molecule due to a steric overlap with Leu29 in CDR H1. **(C)** Ser43 in the light chain of mouse anti-QSOX1 is mutated to Pro in the successful design, likely stabilizing the molecule by rigidifying the loop. Mutation to Ala in the consensus IGKV1 framework likely has little effect. **(D)** Crystal structure of mɑHEWL (gray, PDB entry 1MLC) aligned to CUMAb model of successful design hɑHEWL (blue) and CUMAb model of unsuccessful design hɑHEWL.5 (yellow). Position 34 in the light chain CDR L1 makes two key hydrogen bonds to the side chain of Ser91 in CDR L3 and to the backbone amide of Tyr50 in L2. Because this residue is not in the interface with the antigen, it is mutated in SDR grafting even though it is in a CDR. In the consensus framework IGKV1, this position is mutated to Ala, likely destabilizing CDRs L2 and L3 due to the loss of the hydrogen bonds. Strikingly, in successful CUMAb design 1, this position is mutated to Asn, which is likely to retain the hydrogen bonds.

## Discussion

Instead of using only a few dozen homologous frameworks as in the majority of conventional humanization strategies, CUMAb uses all combinations of possible human gene segments (>20,000 for each antibody) and ranks them by energy. Although genes belonging to a single subgroup are homologous, they contain mutations, including in vernier positions that may stabilize the specific CDRs of the parental antibody. Structural analysis shows that low-energy designs retain critical framework-CDR interactions that may be eliminated in homology-based humanization. Consequently, the lowest-energy designs from this large space of possible frameworks are more likely than the highest homology ones to retain not only binding affinity, but also retain or increase stability and expressibility relative to the parental antibody. Therefore, CUMAb may offer a one-shot strategy to improve stability and expressibility, even for human-sourced antibodies, while maintaining or increasing humanness.

Crucially, CUMAb eliminates the need for back mutations even when starting from an Fv modeled structure. Thus, this approach increases the speed and scope of antibody humanization to potentially any animal antibody without the traditional bottlenecks of obtaining crystallographic structures to enable accurate structure-based humanization^50^. Furthermore, CUMAb is automated and, in most of the cases presented here, required experimental screening of only a handful of constructs, substantially reducing time and cost. It may therefore be applied at scale to dozens of antibodies in parallel. In humanizing the QSOX1 targeting antibody, we tested 15 constructs and found several unique alternatives that exhibited stability and binding properties on par with those of the corresponding animal antibody. Because these alternative designs differ by dozens of surface mutations, they may exhibit differences in critical developability properties, such as nonspecificity and colloidal stability^28, 29, 51^. Further investigation is required to verify the reliability and generality of applying CUMAb to address developability challenges. Since the CUMAb mutations are limited to the framework, the approach is compatible with other design methods that mutate the CDRs to improve aggregation resistance^52, 53^. Additionally, in the current study, the antibody CDRH3 segments were 7-13 amino acids long (Kabat numbering), a size range that is typical of mouse antibodies^54^ and post-phase-1 clinical-stage antibody therapeutics^55^. Longer CDRH3s carry more uncertainty in modeling and design, and it remains to be seen whether they can be humanized reliably using CUMAb.

Finally, we note that CUMAb is based on the fundamental insight that antibody stability and activity are determined by the entire Fv, including the framework positions on which the CDRs rest^16, 56^ and the interfaces between the light and the heavy chains^14, 57^. This insight may help address other important challenges in antibody engineering, leading to general, reliable, and rapid antibody design strategies.

## Data Availability

Amino acid sequences of Fvs for QSOX1, AXL, and HEWL targeting antibodies are available in Supplementary Data Set 3. Human antibody germline sequences were taken from the IMGT reference database (https://www.imgt.org/vquest/refseqh.html) and sequences for FDA-approved humanized antibodies were taken from the TheraSAbDab database (http://opig.stats.ox.ac.uk/webapps/newsabdab/therasabdab/search). The crystal structure of hɑQSOX1.4 is available through Protein Data Bank (PDB) accession ID 8AON.

## Code Availability

RosettaScripts^58^ and command lines for finding the interface residues between the antibody Fv and antigen (Interface.xml, Interface_command_line.txt), relaxing the antibody Fv (Relax.xml, Relax_command_line.txt), and running CUMAb atomistic calculations (CUMAb.xml, CUMAb_command_line.txt) are available as supplemental files. The corresponding flag file (flags.txt) is provided as a supplemental file.

## Acknowledgments

We thank Adva Mechaly for critical reading and Ron Diskin and Olga Khersonsky for advice. Research in the Fleishman lab was supported by the European Research Council through a Consolidator Award (815379), the Dr. Barry Sherman Institute for Medicinal Chemistry, and a donation in memory of Sam Switzer. Research in the Fass lab was supported by the European Research Council through a Proof of Concept grant (825076). Research in the Tessier lab was supported by the National Institutes of Health (RF1AG059723 and R35GM136300) and National Science Foundation (1804313). We acknowledge the European Synchrotron Radiation Facility for the provision of beam time on ID30B, and we thank Andrew McCarthy for his assistance. The collaboration between the Yarden and Fleishman labs was supported by the Weizmann Institute’s BINA framework.

## Author contributions

AT and SJF wrote the manuscript with input from all authors. AT developed the humanization algorithm and designed all humanized sequences. LK performed experimental work related to anti-QSOX1 antibodies with the exception of the developability experiments, LK, NY, and DF performed crystallography data collection, and LK and DF analyzed the data. EM performed developability experiments for anti-QSOX1 antibodies, EM and PT analyzed the data. AN performed IP assay for anti-AXL antibody. ML and IZ expressed and purified anti-AXL antibodies. RK performed experimental work related to anti-MUC16 antibodies, and RK and JA analyzed the data. AT and LK expressed and purified anti-HEWL antibodies. AT, YFS, and YGW performed nano-DSF and SPR characterization of anti-AXL antibodies and analyzed the data together with SJF, ML, and YY. AT, YSF, and YGW performed nano-DSF and SPR characterization of anti-HEWL antibodies and analyzed the data together with SJF.

## Methods

### Computational Methods

All Rosetta design simulations used git version d9d4d5dd3fd516db1ad41b302d147ca0ccd78abd of the Rosetta biomolecular modeling software, which is freely available to academics at http://www.rosettacommons.org.

### A database of antibody germline sequences

Antibody germline sequences were retrieved from the IMGT reference database^59^ (downloaded July 29, 2020). For each gene, only the first allele that was annotated as functional was taken. Partial or reverse-complementary genes were pruned. If allele one contains more than two cysteines, a different allele was taken with two cysteines if available. This resulted in 54 heavy chain V gene sequences, six heavy chain J gene sequences, 39 light chain kappa V gene sequences, five light chain kappa J gene sequences, 30 light chain lambda V gene sequences, and five light chain kappa J gene sequences. We then combined these genes all-against-all (V and J for both light and heavy chains) for kappa and lambda light chains separately, resulting in 63,180 sequences for kappa light chains and 48,600 for lambda light chains.

### Computational CDR grafting

We defined the CDRs once for all the antibody targets in this study based on previous definitions^20, 21^ and visual inspection of antibody crystal structures. Starting from a molecular structure of the parental antibody, we use HMMer to identify the sequence segments corresponding to the variable region and classify the light chain as kappa or lambda^60^. For each germline sequence corresponding to the light chain classification, the CDRs of the germline sequence are replaced with the CDRs of the parental (in the current study, mouse) antibody. Any sequence that contains an Asn-Gly or Asn-X-Ser/Thr (where X is not Pro) outside of the CDRs is removed. This typically results in >20,000 unique sequences per parental antibody.

### Energy ranking of CDR-grafted sequences

If provided a crystal structure, as a first step, the structure of the parental antibody is relaxed through cycles of sidechain and harmonically restrained backbone minimization and combinatorial sidechain packing in the entire Fv (see Relax.xml RosettaScript^58^). If a bound structure is provided, residues in the interface between the Fv and the antigen are identified using Rosetta and held fixed during the relaxation (see Interface.xml). Additionally, if a bound structure is provided, the entire antibody Fv-antigen complex is relaxed.

Each CDR-grafted germline sequence is then threaded onto the relaxed Fv structure and relaxed in the same manner with fewer cycles (see CUMAb.xml). Sequences are ranked according to the all-atom energy ref2015^23^. Any model that exhibits a Cɑ-carbonyl O RMSD greater than or equal to 0.5 Å in any of the CDRs is excluded from further consideration. This threshold was selected by visual inspection of designs and is intended to limit conformational changes that may impact the structure of the CDRs. Sequences are clustered according to V gene subgroup as defined by IMGT, meaning that we take only one sequence from each V gene combination. Sequences were visually inspected, and, in some cases, the highest ranking representative for a cluster was replaced with a slightly lower ranking one in order to re-use sequences in different clusters and thus minimize cloning.

When starting from a model rather than an experimental structure, the pipeline is identical, except that sequences are not excluded for deviating from the model structure, thus allowing for modeling inaccuracies. This will allow some flexibility to make small changes in the backbone conformations of the CDR loops, which may not be modeled completely accurately by AlphaFold, particularly CDR H3.

### Model structures of antibodies

All model structures were generated using the ColabFold^40^ (in multimer setting^41^) with default settings. Top ranked model by pTM score was chosen for design. In the specific case of anti-mucin16 Ab2, we noticed that the AlphaFold model exhibited an unrealistic strand swapping between the light and heavy chains. We therefore used AbPredict^38^, visually inspected the three suggested models and selected the second as the starting point for design.

### Computational SDR grafting

The parental antibody is classified as having a kappa or lambda light chain as described above. We use Rosetta to identify residues in the interface between the antibody Fv and the antigen (see interface.xml). Full acceptor frameworks are assembled by combining each human germline heavy V, heavy J, light V, and light J. These acceptor frameworks are thus nearly complete germline antibodies that are only missing D genes in CDR H3. Acceptor frameworks are then selected using the following criteria: The frameworks must have the same CDR length in all CDRs as the parental antibody, excluding CDR H3. Additionally, the frameworks must have an H3 length that is equal to or shorter than the H3 length of the parental antibody. If the H3 length of the parental framework is shorter than that of the parental antibody, a number of residues equal to the difference in length of the two H3s from the parental antibody are inserted into the germline sequence. The acceptor frameworks are then threaded and relaxed as described above. Sequences are clustered according to V gene and heavy J gene subgroup.

### Evaluation of humanness of antibody sequences

Antibody heavy or light chain sequences are submitted to the IgBLAST^61^ web server, and the sequence identity to the top hit is taken. For additional verification, antibody sequences are submitted to the Hu-mAb^26^ and BioPhi^27^ web servers using default settings except for that BioPhi the prevalence threshold is set to medium.

### Generation of surface electrostatic charge maps for anti-QSOX1 antibodies

Charge maps were generated using the APBS Electrostatic plug-in^62^ in PyMOL version 2.4.0 using default settings, with the exception of the Molecular Surface Visualization range, which was set to +/- 3.50.

### Sequence alignments of antibodies to highest homology human V gene germlines

IgBlast webserver^61^ was used to identify the human V gene germlines with the highest homolgy to the parental antibody for both the light and heavy chain (top-hit was selected). This combination of human germline genes was aligned to the successful CUMAb designs using the Clustal Omega webserver^63^ and plots were done in Jalview^64^ (Version: 2.11.2.5).

## Experimental Methods

### Expression and purification of anti-human QSOX1 antibodies

Expression plasmids for CUMAb light and heavy chains were constructed on the basis of the AbLIFT18 chimeric antibody^14^ using restriction-free procedures to replace the variable regions. Plasmids were purified from the *E. coli* XL-1 strain. Proteins were produced by transient transfection of plasmids into HEK293F cells. Transfection was done using the PEI Max reagent (Polysciences Inc.) with a 1:3 ratio (w/w) of DNA to PEI at a concentration of 1 million cells per ml. Co-transfections were done with a 1:1 ratio of light- and heavy-chain DNA, while the total amount of DNA remained the same: 1 µg DNA per ml HEK culture. Six days after transfection, the culture medium was collected and centrifuged for 10 min at 500 g to pellet cells. The supernatant was then centrifuged for 10 min at 3200 g to pellet any remaining particles. The supernatant was filtered through a 0.22 µm filter, and the antibodies were purified by protein G affinity chromatography (GE Healthcare). Antibodies were then dialyzed into PBS pH 8 overnight.

### Expression and purification of human QSOX1 proteins

Construction of pET15b expression vectors for human QSOX1, used for activity assays (residues 34–546 of HsQSOX1), and of its oxidoreductase fragment (QSOX1trx, residues 36-275 of HsQSOX1), used for co-crystallization with a Fab fragment of a CUMAb design, were previously described^65^. Both include an amino-terminal His_6_ tag. The His_6_ tag of QSOX1trx is cleavable by a downstream thrombin cleavage site and was removed by thrombin cleavage prior to crystallization. QSOX1 proteins were purified by nickel-nitrilotriacetic acid (Ni-NTA) chromatography (GE Healthcare).

### QSOX1 activity/inhibition assays

#### QSOX1 activity on Zymogen granule membrane protein 16 (ZG16) for initial antibody screening

ZG16 contains two cysteines in a flexible, accessible CXXC motif that can be oxidized in vitro by QSOX1^66^. ZG16 fused to maltose binding protein (MBP, ∼45 kDa, no cysteines) was used to assay QSOX1 activity in the presence of supernatants from antibody-producing HEK293F cells. Prior to the assay, MBP-ZG16 was reduced by incubation with DTT, excess DTT was removed on a PD-10 column (GE Healthcare), and reactions of the reduced MBP-ZG16 and QSOX1 (50 nM) were initiated in the presence of the antibody variants, supplied by adding clarified HEK293F culture supernatants diluted into the reaction mixture. The mixture contained 8 microliters HEK293F supernatant, 2.8 microliters reduced MBP-ZG16 (2.6 micromolar final concentration), and 1.2 microliters QSOX1 (50 nM final concentration). The reactions were stopped by adding maleimide-functionalized polyethylene glycol of molecular weight 5000 Da (PEG-mal 5000, Iris biotech), which covalently modifies any residual free thiols, resulting in slower migration in SDS-PAGE. Loading buffer was added to the reaction mixture and the whole mixture was run on an SDS-PAGE gel. Increased amounts of modified MBP-ZG16 relative to the uninhibited reaction indicated inhibition in this screen.

#### Plasma-QSOX1 inhibition

All procedures involving human samples were approved by the Weizmann Institute of Science Institutional Review Board. Five cc of blood was collected by venipuncture from healthy human donors with informed consent. Heparin was used as an anticoagulant, and blood was centrifuged for 15 min at 300 g. The plasma was collected and used as a native QSOX1 source in QSOX1 inhibition reactions conducted in 100 μl reactions as follows: 10 μl plasma was incubated at 37° C with 25 nM of the CUMAb antibody hɑQSOX1.1 in 50 mM potassium phosphate buffer, pH 7.5, 65 mM NaCl, and 1 mM EDTA. The reaction was initiated by the addition of dithiothreitol (DTT) to a final concentration of 300 µM. After 30 min at room temperature, 50 μl were removed and added to 725 μl of the above buffer and 25 μl of 2 mM 5,5′-dithiobis-(2-nitrobenzoic acid) (DTNB), which reacts with residual DTT and produces the chromogenic product 5-thio-2-nitrobenzoate (TNB) with absorption at 412 nm.

### QSOX1 inhibition constant determination

A Clark-type oxygen electrode (Hansatech Instruments) was used to monitor changes in dissolved oxygen concentration as a measure of QSOX1 activity. QSOX1 (25 nM) and various concentrations (1-100 nM) of purified antibody were assayed in 50 mM potassium phosphate buffer, pH 7.5, 65 mM NaCl and 1 mM EDTA. Reactions were started by injection of DTT to a concentration of 200 μM in the electrode reaction chamber.

Two independent progress curves were collected for each antibody concentration, and initial slopes were calculated. The background decrease in oxygen concentration due to the presence of DTT and antibody was measured and subtracted from the initial slopes to obtain the rates of QSOX1 activity in the presence of various concentrations of each antibody. The ratios of the initial rates of QSOX1 in the presence and absence of inhibitor were plotted as a function of inhibitor concentration. The data were fit to the Morrison equation for a tight binding inhibitor (GraphPad Prism version 8.4.3 for Windows, GraphPad Software, San Diego, California USA, www.graphpad.com) to determine the inhibition constants of the antibodies.

### Charge-stabilized self-interaction nanoparticle spectroscopy (CS-SINS) measurements

CS-SINS was measured as reported previously^28^. Briefly, capture antibody (0.8 mg/mL; goat anti-human Fc, Jackson ImmunoResearch, 109-005-008) and polylysine (2.67 mg/mL; ≥70 kDa, Fisher Scientific, ICN19454405) were mixed at a ratio of 90%:10% w/w. Gold nanoparticles (Ted Pella, 15705-1) were transferred to 8 tubes (USA Scientific, 1615-5500), sedimented at 21630 xg, and 95% of the supernatant was removed (e.g., 1150 out of 1200 μL) and discarded. The capture antibody and polylysine mixture was immobilized on concentrated gold nanoparticles (equal parts of capture antibody/polylysine mix and concentrated gold) and incubated overnight at room temperature. Conjugates (5 μL) were then incubated with dilute antibody solutions (11.1 μg/mL, 45 μL) for 4 h at room temperature. Absorbance spectra were measured on a Biotek Synergy Neo plate reader (Biotek, Winooski, VT) in 1 nm increments between 450 and 650 nm. A quadratic equation was fit to describe the forty data points surrounding the maximum measured absorbance. The inflection point of the quadratic was calculated to determine the plasmon wavelength. Plasmon wavelengths were normalized between two fit parameters and calibrated against a panel of five antibodies (tocilizumab, cetuximab, evolocumab, denosumab, and omalizumab) run with each experiment.

### PolySpecificity Particle (PSP) measurements

Non-specific binding analysis was performed as reported previously^29^. Briefly, Protein A Dynabeads (Invitrogen, 10002D) were washed three times with PBS with 1 mg/mL BSA (PBSB). The Dynabeads (1.6 μg) were incubated with antibodies (85 μL, 15 g/mL) overnight at 4 °C. The coated beads were then washed twice by centrifugation (3500x g for 4 min) with PBSB. Biotinylated soluble membrane proteins (SMP) from CHO cells (0.1 mg/mL), prepared as described previously^30^, were incubated with the washed beads at 4 °C for 20 min. The beads were then washed once (PBSB) and incubated with 0.001x streptavidin-AF647 (Invitrogen, S32357) and 0.001x goat anti-human Fc F(ab’)2 AF488 (Invitrogen, H10120) on ice for 4 min. The beads were washed, resuspended in PBSB, and analyzed via flow cytometry. Results were normalized between emibetuzumab and elotuzumab as the high and low binding controls, respectively. The results were adjusted to the same scale as previously reported^29^, which was normalized between ixekizumab and elotuzumab.

### Purification and crystallization of the hQSOX1trx/hαQSOX1.4 complex

Purified QSOX1trx was concentrated to 10 mg/ml. The purified CUMAb antibody hαQSOX1.4 was concentrated to 1.5 mg/mL in PBS and digested at 37 °C with papain (Sigma) at a 1:20 papain:hαQSOX1.4 molar ratio. Prior to use, papain was activated by dissolving in PBS supplemented with 20 mM EDTA and 20 mM cysteine. Digestion was stopped after 4 h using leupeptin, and the digested antibody was dialyzed against PBS, pH 8. The Fab fragment of hαQSOX1.4 was purified by protein A affinity chromatography (Cytiva). Purified hαQSOX1.4 Fab was incubated for 1 h at 4 °C with a 2- to 3-fold excess of QSOX1trx, and the complex was isolated by size-exclusion chromatography in 10 mM Tris, pH 7.5, and 100 mM NaCl. The complex was concentrated using a centrifugal concentrator to 10 mg/mL. Crystals were grown by hanging-drop vapor diffusion at 293 K by mixing 1 μl of protein complex solution with 1 μl of well solution and suspending over well solution. Initial branched, feather-like crystals were observed when well conditions were 14% w/v polyethylene glycol 4 kDa and 0.1 M ammonium phosphate dibasic. These crystals were seeded into drops in similar conditions (11-14% w/v polyethylene glycol 4 kDa and 0.05-0.15 M ammonium phosphate dibasic) to produce individual crystal rods. These crystals were soaked in a similar solution with an addition of 20% glycerol and flash frozen in liquid nitrogen.

### X-ray data collection and model building

X-ray data were collected at a wavelength of 0.9763 Å on the ID30B beamline at the European Synchrotron Radiation Facility, using a helical collection mode. Molecular replacement was performed using Phenix^67^ with the hQSOX1trx structure from PDB entry 4IJ3 and an Alphafold prediction model for the hαQSOX1.4 Fab structure as input models. The initial complex model was then iteratively rebuilt and refined using Coot^68^ and Phenix^67^, respectively. The refined structure was deposited to the PDB with entry code 8AON.

### Secreted Fab (anti-HEWL) expression and purification

The variable regions of the different heavy and light chains were cloned separately, upstream of IgG1 human Ab Fab scaffolds, into p3BNC plasmids. Antibody Fabs were produced by transient transfection of plasmids into HEK293F cells. Transfection was done using the PEI Max reagent (Polysciences Inc.) with a 1:3 ratio (w/w) of DNA to PEI at a concentration of 1 million cells per ml. Co-transfections were done with a 1:1 ratio of light- and heavy-chain DNA, while the total amount of DNA remained the same: 1 µg DNA per ml HEK culture. Six days after transfection, the culture medium was collected and centrifuged for 10 min at 500 g to pellet cells. The supernatant was then centrifuged for 10 min at 3200 g to pellet any remaining particles. The supernatant was filtered through a 0.22 µm filter, and the antibodies were purified by nickel-nitrilotriacetic acid (Ni-NTA) chromatography (GE Healthcare). Antibodies were then dialyzed into PBS pH8 overnight.

### Expression and purification of humanized anti-human AXL

Anti-hAXL was generated as a human IgG1/IgK subclass. The sequences of the variable regions of the heavy and light chains were synthesized by Twist Bioscience, polymerase chain reaction (PCR)–amplified, and cloned into mammalian expression vectors (purchased from Oxford genetics) in frame with the constant regions of the human IgG1 heavy chain or human kappa light chain, respectively. Plasmid sequences were validated by direct sequencing (Life Sciences Core Facility, Weizmann Institute of Science). To produce antibodies, antibody heavy and light chain expression vectors were transiently transfected into Expi293 cells (Thermo Fisher Scientific). The secreted antibodies in the cell supernatant were purified using protein G Sepharose 4 Fast Flow Resin (GE Healthcare). Purified antibodies were dialyzed in phosphate-buffered saline and sterile-filtered (0.22 μm). Purity was assessed by SDS-PAGE and Coomassie staining and was estimated to be >90% (data not shown).

### Immunoprecipitation of anti-AXL antibodies

Immunoprecipitation assays used protein G beads. Following gel electrophoresis, proteins were transferred onto a nitrocellulose membrane. The membrane was blocked in TBST buffer (0.02 M Tris-HCl (pH 7.5), 0.15 M NaCl, and 0.05% Tween 20) containing albumin (3%), blotted with a primary antibody overnight at 4° C, washed in TBST and incubated for 30 min with a secondary antibody conjugated to horseradish peroxidase.

### Apparent Tm measurements of anti-HEWL and anti-AXL antibodies

The apparent melting temperature of the antibodies was determined by the Prometheus NT. 48 instrument (NanoTemper Technologies). Full length IgGs (anti-AXL) or Fabs (anti-HEWL) were diluted to 0.5 mg/ml (anti-AXL) or 0.1 mg/mL (anti-HEWL) in 1X PBS. The temperature was ramped from 20°C to 95°C at 1.0°C/min, and Tm was measured. Calculations were done using Prometheus NT. 48 default software. All antibodies were repeated.

### Surface-plasmon resonance of anti-AXL antibodies

Surface plasmon resonance (SPR) experiments on the anti-AXL antibodies were carried out on a Biacore S200 instrument (GE Healthcare, Sweden) at 25°C with PBST (1X PBS + 0.05% Tween-20). For binding analysis, approximately 1,000 response units (RU) of AXL (R&D systems) were conjugated to a CM5 sensor chip using NHS/EDC chemistry. Samples of different antibody concentrations were injected over the surface at a flow rate of 30 μL/min for 240 s, and the chip was washed with buffer for 180 s. After each injection, surface regeneration was performed with a 30 s injection of 1 mM NaOH at a flow rate of 30 uL/min. The first channel contained no immobilized ligand, and its sensogram was subtracted from the channel used to measure binding. The acquired sensograms were analyzed using the device’s software version 1.1, and kinetic fits to 1:1 binding model were performed.

### Expression and purification of anti-MUC16 antibodies

The heavy and light chain variable regions of cɑMUC16_Ab1, hɑMUC16_Ab1, cɑMUC16_Ab2, hɑMUC16_Ab2.1 and hɑMUC16_Ab2.2 were ordered as synthetic gBlocks Gene Fragments (IDT) and subcloned into DNA expression plasmids containing the constant domain of human IgG1 (PSF-CMV-HUIGG1 HC OGS527, Sigma) or human Kappa (PSF-CMV-HUKAPPA LC OGS528, Sigma), respectively. Plasmids were purified from the *E. coli* DH5-alpha strain using ZymoPURE II Plasmid Midiprep Kit (Zymo Research). Recombinant antibodies were produced by transient transfection of plasmids into HEK293 cells pre-plated in 100 mm culture dishes with 75-85% confluency. Transfection was done using PEI 25K (Polysciences Inc.) with a 1:3 ratio (w/w) of DNA to PEI and a 1:1.5 ratio of heavy- and light-chain DNA, while the total amount of DNA remained the same: 10 µg DNA per one culture plate. Three days after transfection, the culture medium was collected and centrifuged for 10 min at 3300xg to pellet cells and debris. The supernatant was then filtered through a 0.45 µm filter, and the antibodies were purified by protein G beads (Thermo Fisher Scientific).

### Measurement of anti-MUC16 antibody expression levels

Anti-MUC16 antibodies were expressed as described before. A portion of the supernatant was used to determine antibody concentration by ELISA. Serial dilution was performed for each antibody supernatant and samples were coated in Nunc MaxiSorp™ flat-bottom ELISA plates (Thermo Fisher Scientific) overnight at 4 ℃. After blocking for 1 hour at RT with 5% FBS, the plates were washed and incubated with anti-human IgG antibody conjugated to HRP (Jackson ImmunoResearch Laboratories, Inc.) for 1 hour at RT. The plates were then washed and developed for 2.5 min. with 3,3′,5,5′-Tetramethylbenzidine (TMB) (Sigma) before stopping the reaction with sulfuric acid. The plates were read at 450 nm using Epoch Microplate Spectrophotometer (Agilent BioTek). An antibody with a pre-determined concentration was used to construct a standard curve and calculate anti-MUC16 antibody expression levels.

### Measurement of anti-MUC16 antibody affinity to MUC16 using ELISA

Serial dilutions of recombinant MUC16 protein were performed and samples ranging from 100 nM to 0.195 nM were coated in Nunc MaxiSorp™ flat-bottom ELISA plates (Thermo Fisher Scientific) overnight at 4 ℃. After blocking for 1.5 hours at RT with 5% FBS, the plates were washed and incubated with anti-MUC16 antibodies in a concentration of 7 nM for 1h at RT. Afterwards, the plates were washed and incubated with anti-human IgG antibody conjugated to HRP (Jackson ImmunoResearch Laboratories, Inc.) for 1 hour at RT. The plates were then washed and developed for 1 min. with 3,3′,5,5′-Tetramethylbenzidine (TMB) (Sigma) before stopping the reaction with sulfuric acid. The plates were read at 450 nm using Epoch Microplate Spectrophotometer (Agilent BioTek). After subtracting the background of the primary antibodies to the blocking reagent or the background of the secondary antibody to the recombinant MUC16 (whichever was higher), a non-linear curve fitting to the one site, specific-binding equation was performed using GraphPad Prism 9 (Version 9.4.1 (458)) to estimate the Kd of each antibody.

### FACS analysis of anti-MUC16 antibody binding to MUC16

The MUC16-expressing ovarian cancer cell line, OVCAR-3, was used to measure antibody binding to natively expressed MUC16. The cells were harvested with trypsin, washed with PBS and stained with a viability dye in MACS buffer for 10 min. at 4 ℃. After washing with MACS buffer, the cells were fixed with acetone for 1h at 4 ℃. Then, the cells were washed twice with PBS and fixed with 10% goat serum in PBS with 0.3 M glycine for 45 min. at RT, followed by one wash and then stained with anti-MUC16 antibodies for 1h at 4 ℃ in a final antibody concentration of 3.33 ug/ml in MACS buffer. Afterwards, the cells were washed and stained with goat anti-Human IgG Fc Secondary Antibody PE 1:200 (eBioscience™) for 45 min. at 4 ℃. After washing with MACS buffer the samples were analyzed by BD FACSCanto II (BD biosciences) after pregating on live single cells for FSC-area vs. FSC-height. Alternatively, HEK293 cells were transfected with a DNA plasmid encoding a truncated MUC16 containing the predicted epitope of the anti-MUC16 antibodies. The cells were kept alive, and except from fixation, the cells were prepared and analyzed in the same manner as OVCAR-3 cells. The data were analyzed by Flowjo (v10, BD biosciences).

### Yeast surface display of anti-HEWL antibodies

Yeast-display experiments were conducted essentially as described^48^. Briefly, yeast cells were grown in selective medium SDCAA overnight at 30°C. The cells were then resuspended in 1 ml of induction medium and incubated at 20°C for 20 h. Roughly 10^7^ cells were then used for yeast-cell surface display experiments: cells were subjected to primary antibody (mouse monoclonal IgG1 anti-c-Myc (9E10) sc-40, Santa Cruz Biotechnology) for expression monitoring and biotinylated ligand at 90 nM lysozyme (GeneTex) in PBS-F for 30 min at room temperature. The cells then underwent a second staining with fluorescently labeled secondary antibody (AlexaFluor488—goat-anti-mouse IgG1 (Life Technologies) for scFv labeling, Streptavidin-APC (SouthernBiotech) for ligand labeling) for 10 min at 4°C. Next, the cell fluorescence was measured. Data was analyzed using FlowJo (v10, BD biosciences).

### Surface-plasmon resonance of anti-HEWL antibodies

Surface plasmon resonance experiments on the anti-HEWL Fabs were carried out on a Biacore S200 instrument (GE Healthcare, Sweden) at 25°C with HBS-N EP+ [10 mM Hepes, 150 mM NaCl, 3 mM EDTA, 0.005% vol/vol surfactant P20 (pH 7.4)]. For binding analysis, approximately 300 response units (RU) of Fab were conjugated to a CM5 sensor chip using NHS/EDC chemistry. Samples of different HEWL (Sigma) concentrations were injected over the surface at a flow rate of 30 μL/min for 120 s, and the chip was washed with buffer for 280 s (mouse Fab) or 130 s (humanized Fab). The first flow cell contained no Fab and was subtracted from the channel used to measure binding as a reference. The acquired data were analyzed using the device’s software, and kinetic fits to 1:1 binding model were performed.

## Supplement

**Supplementary Table 1:**
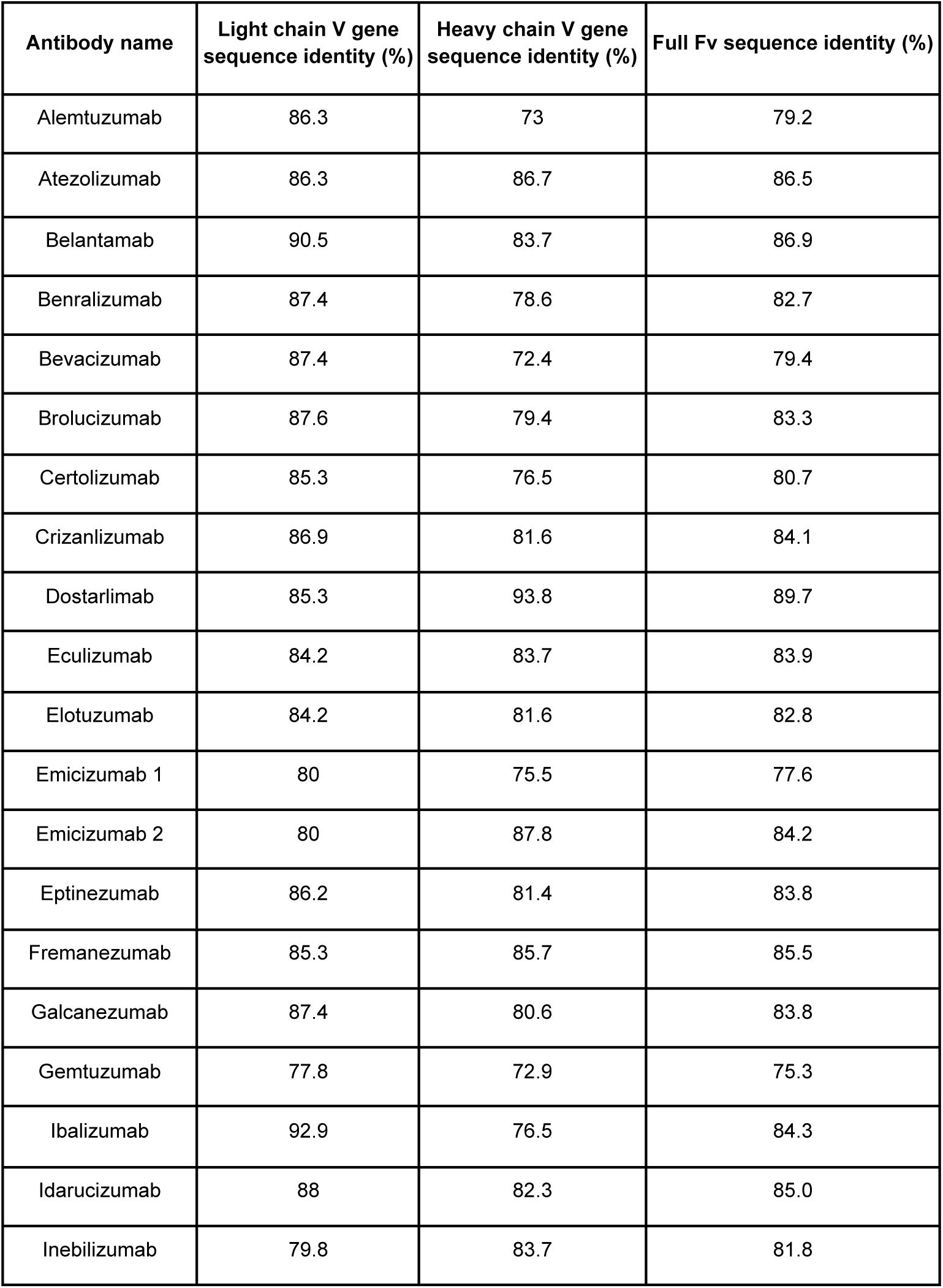
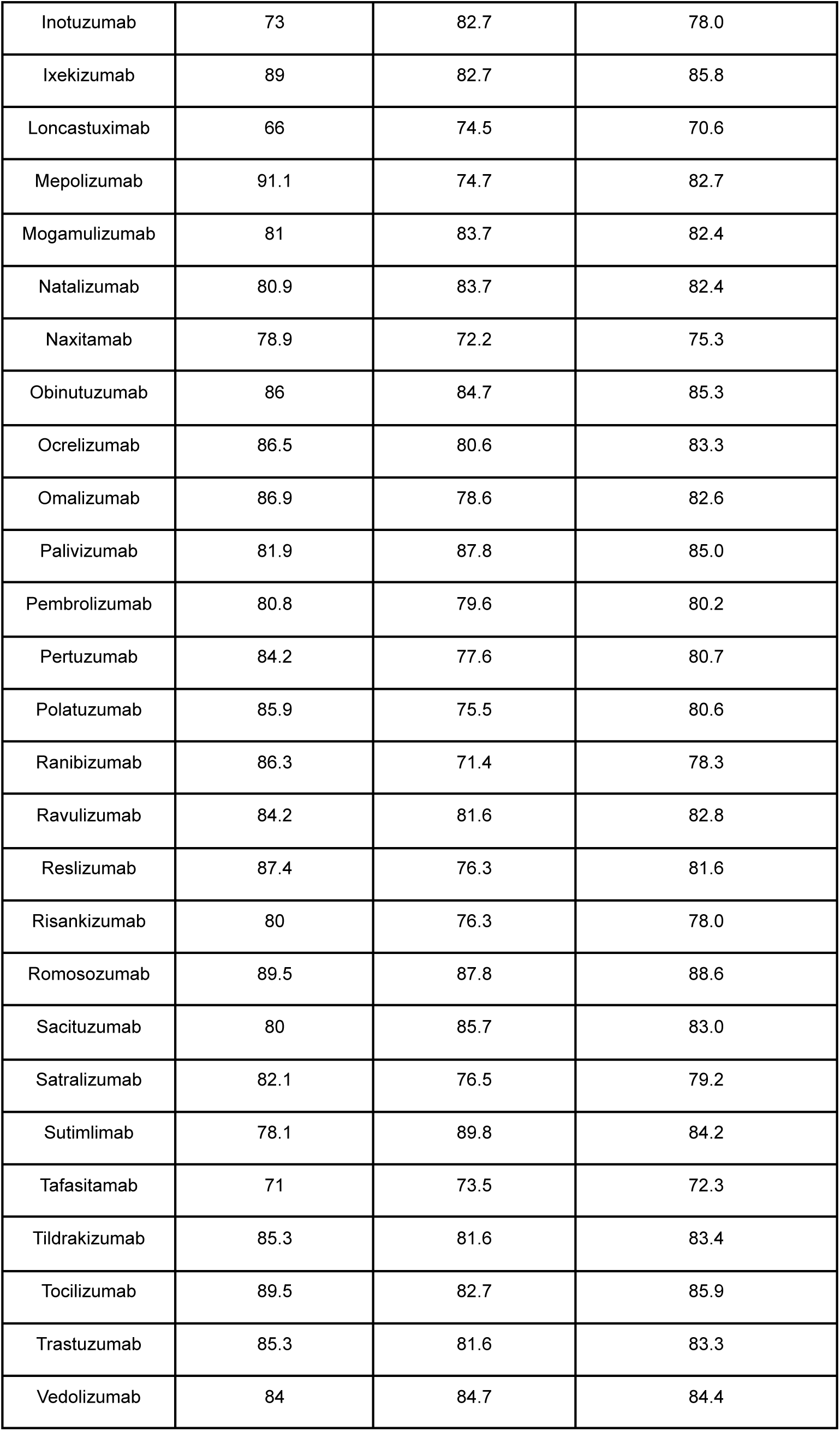
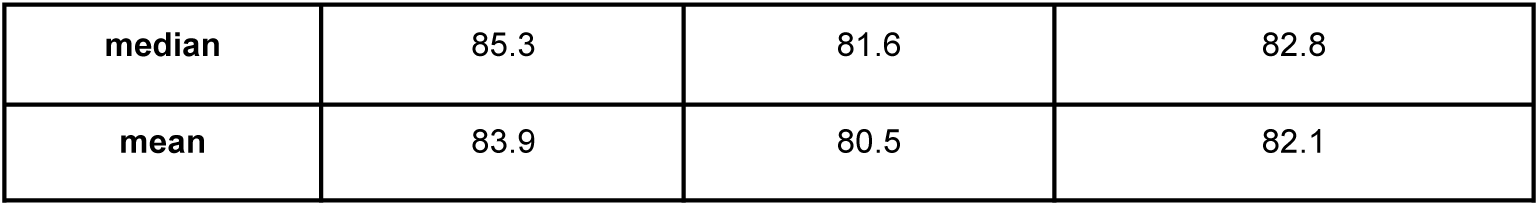
Sequence identity to the human germline of 47 FDA approved humanized antibodies^1^. % identity computed using IgBLAST^2^. Antibody sequences were taken from Thera-SAbDab^3^.

| Antibody name | Light chain V gene sequence identity (%) | Heavy chain V gene sequence identity (%) | Full Fv sequence identity (%) |
| --- | --- | --- | --- |
| Alemtuzumab | 86.3 | 73 | 79.2 |
| Atezolizumab | 86.3 | 86.7 | 86.5 |
| Belantamab | 90.5 | 83.7 | 86.9 |
| Benralizumab | 87.4 | 78.6 | 82.7 |
| Bevacizumab | 87.4 | 72.4 | 79.4 |
| Brolucizumab | 87.6 | 79.4 | 83.3 |
| Certolizumab | 85.3 | 76.5 | 80.7 |
| Crizanlizumab | 86.9 | 81.6 | 84.1 |
| Dostarlimab | 85.3 | 93.8 | 89.7 |
| Eculizumab | 84.2 | 83.7 | 83.9 |
| Elotuzumab | 84.2 | 81.6 | 82.8 |
| Emicizumab 1 | 80 | 75.5 | 77.6 |
| Emicizumab 2 | 80 | 87.8 | 84.2 |
| Eptinezumab | 86.2 | 81.4 | 83.8 |
| Fremanezumab | 85.3 | 85.7 | 85.5 |
| Galcanezumab | 87.4 | 80.6 | 83.8 |
| Gemtuzumab | 77.8 | 72.9 | 75.3 |
| Ibalizumab | 92.9 | 76.5 | 84.3 |
| Idarucizumab | 88 | 82.3 | 85.0 |
| Inebilizumab | 79.8 | 83.7 | 81.8 |
| Inotuzumab | 73 | 82.7 | 78.0 |
| Ixekizumab | 89 | 82.7 | 85.8 |
| Loncastuximab | 66 | 74.5 | 70.6 |
| Mepolizumab | 91.1 | 74.7 | 82.7 |
| Mogamulizumab | 81 | 83.7 | 82.4 |
| Natalizumab | 80.9 | 83.7 | 82.4 |
| Naxitamab | 78.9 | 72.2 | 75.3 |
| Obinutuzumab | 86 | 84.7 | 85.3 |
| Ocrelizumab | 86.5 | 80.6 | 83.3 |
| Omalizumab | 86.9 | 78.6 | 82.6 |
| Palivizumab | 81.9 | 87.8 | 85.0 |
| Pembrolizumab | 80.8 | 79.6 | 80.2 |
| Pertuzumab | 84.2 | 77.6 | 80.7 |
| Polatuzumab | 85.9 | 75.5 | 80.6 |
| Ranibizumab | 86.3 | 71.4 | 78.3 |
| Ravulizumab | 84.2 | 81.6 | 82.8 |
| Reslizumab | 87.4 | 76.3 | 81.6 |
| Risankizumab | 80 | 76.3 | 78.0 |
| Romosozumab | 89.5 | 87.8 | 88.6 |
| Sacituzumab | 80 | 85.7 | 83.0 |
| Satralizumab | 82.1 | 76.5 | 79.2 |
| Sutimlimab | 78.1 | 89.8 | 84.2 |
| Tafasitamab | 71 | 73.5 | 72.3 |
| Tildrakizumab | 85.3 | 81.6 | 83.4 |
| Tocilizumab | 89.5 | 82.7 | 85.9 |
| Trastuzumab | 85.3 | 81.6 | 83.3 |
| Vedolizumab | 84 | 84.7 | 84.4 |
| **median** | 85.3 | 81.6 | 82.8 |
| **mean** | 83.9 | 80.5 | 82.1 |

**Supplementary Table 2.** CUMAb CDR definitions compared to Kabat and AHO schemes^4, 5^. Position numbering according to Kabat.

|  | **L1** | **L2** | **L3** | **H1** | **H2** | **H3** |
| --- | --- | --- | --- | --- | --- | --- |
| **CUMAb** | 24-34 | 46-55 | 89-97 | 24-35B | 47-58 | 93-102 |
| **Kabat** | 24-34 | 50-56 | 89-97 | 31-35B | 50-65 | 95-102 |
| **AHO** | 25-32 | 50-61 | 91-96 | 24-35B | 51-66 | 95-101 |

**Supplementary Table 3:** Charge mismatches between anti-QSOX1 CUMAb designs. Each pair of sequences was aligned and the number of times positively charged residues (lysine or arginine) or negatively charged residues (aspartate or glutamate) were present at a given position of one sequence and not the other were counted. For each pair, the count of positive mismatches is shown in the cell colored blue and the count of negative mismatches is shown in the transparent cell. The number of total charges in each sequence is as follows: mɑQSOX1 20 positive, 15 negative; hɑQSOX1.1 17 positive, 18 negative; hɑQSOX1.2 18 positive, 17 negative; hɑQSOX1.3 18 positive, 16 negative.

|  | **maQSOX1** | **haQSOX1.1** | **haQSOX1.2** | **haQSOX1.3** |
| --- | --- | --- | --- | --- |
| **maQSOX1** | 0 | 7 | 6 | 12 |
|  | 0 | 3 | 4 | 3 |
| **haQSOX1.1** | 7 | 0 | 5 | 5 |
|  | 3 | 0 | 3 | 4 |
| **haQSOX1.2** | 6 | 5 | 0 | 10 |
|  | 4 | 3 | 0 | 5 |
| **haQSOX1.3** | 12 | 5 | 10 | 0 |
|  | 3 | 4 | 5 | 0 |

**Supplementary Table 4:** Crystallographic data and refinement statistics for the complex between the Fab fragment of hαQSOX1.4 and QSOX1trx (PDB code 8AON). Values in parentheses are for the highest resolution bins.

| Data collection |  |
| --- | --- |
| Space group | $P2_1$ |
| Cell dimensions |  |
| a, b, c (Å) | 55.58, 98.28, 63.12 |
| $\alpha, \beta, \gamma$ (°) | 90, 101.15, 90 |
| Asymmetric unit | 1 Fab-QSOX1trx complex |
| Resolution (Å) | 49.14 - 2.10 (2.18 - 2.10)* |
| Measured reflections | 102,949 (10,755) |
| Unique reflections | 37,687 (3817) |
| Completeness (%) | 97.0 (98.6) |
| Redundancy | 2.7 (2.8) |
| $\langle I/\sigma I \rangle$ | 6.8 (0.6) |
| $R_{\text{merge}}$ | 0.105 (1.637) |
| $R_{\text{pim}}$ | 0.075 (1.127) |
| Refinement |  |
| Resolution (Å) | 49.14 - 2.10 (2.12 - 2.10) |
| Reflections in working set | 69,556 (2257) |
| Reflections in test set | 7728 (254) |
| $R_{\text{work}}/R_{\text{free}}$ | 0.198/0.259 |
| Number of protein atoms | 5,061 |
| Number of water molecules | 211 |
| Mean B-factor | 46 |
| Root mean square deviations |  |
| Bond length (Å) | 0.007 |
| Bond angle (°) | 0.908 |
| Ramachandran plot |  |
| Favored regions (%) | 97.1 |
| Additional allowed regions (%) | 2.9 |
| Disallowed regions (%) | 0 |

**Supplementary Table 5:** Sequence identity between parental antibodies (first in each block) and designs to the human germline. % identity computed using IgBLAST^2^.

| Antibody | Closest heavy V germline match | Closest light V germline match |
| --- | --- | --- |
| maQSOX1 | IGHV4-4 (57%) | IGKV1-33 (66%) |
| haQSOX1.1 | IGHV3-33 (80%) | IGKV4-1 (81%) |
| haQSOX1.2 | IGHV2-5 (86%) | IGKV4-1 (81%) |
| haQSOX1.3 | IGHV3-49 (81%) | IGKVD-29 (78%) |
| haQSOX1.4 | IGHV3-49 (81%) | IGKV4-1 (81%) |
| Consensus H4-K1 | IGHV4-4 (85%) | IGKV1-NL1 (84%) |
| maAXL | IGHV3-23 (79%) | IGKV1-16 (62%) |
| haAXL | IGHV3-23 (89%) | IGKV1D-13 (84%) |
| caMUC16-1 | IGHV1-46 (62%) | IGKV2-30 (79%) |
| haMUC16-1 | IGHV4-31 (82%) | IGKV2-29 (85%) |
| caMUC16-2 | IGHV1-46 (61%) | IGKV4-1 (81%) |
| haMUC16-2.1 | IGHV1-69-2 (84%) | IGKV2D-29 (86%) |
| haMUC16-1.2 | IGHV1-69-2 (84%) | IGKV1-9 (83%) |
| maHEWL | IGHV1-46 (67%) | IGKV3-15 (64%) |
| haHEWL | IGHV1-69-2 (90%) | IGKV1-39 (96%) |

**Supplementary Figure 1:**
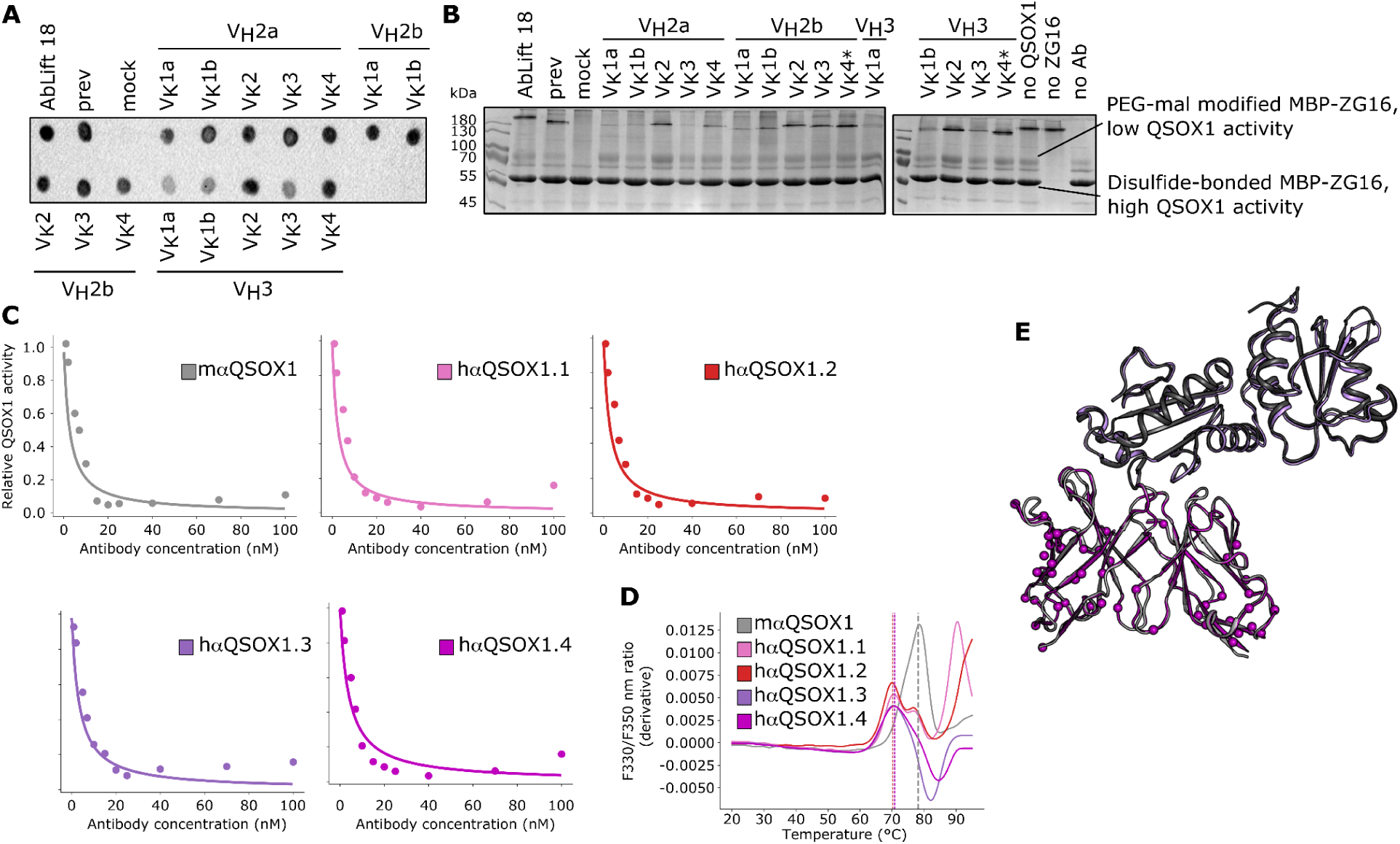
(**A**) Dot-blot analysis of 15 CUMAb designs. 12 designs exhibit comparable expression levels to the AbLIFT18 design of the anti-QSOX1 antibody^6^. CUMAb designs are labeled according to heavy chain V-gene subgroup and then by light chain subgroup according to IMGT. If there are two chains from the same subgroup, they are labeled as “a” or “b” arbitrarily. “Mock” refers to a control transfection where no DNA was added. “Prev” refers to a construct from a prior version of CUMAb. (**B**) Functional screen of 15 CUMAb designs, labeled as in A. Presence of an upper band, corresponding to MBP-ZG16 modified with PEG-mal 5000, indicates remaining unoxidized MBP-ZG16 as a result of inhibited QSOX1 activity. As evident from the “no QSOX1” lane, the MBP-ZG16 used in the experiment was not fully reduced, but the reduced population was sufficient to distinguish between the mock-transfected sample, which lacks antibody, and the antibody-containing samples, which all show some PEG-mal-modified species indicative of QSOX1 inhibition. Designs marked with * were further analyzed. (**C**) Titration of relative QSOX1 activity at different antibody concentrations for mɑQSOX1 and four CUMAb designs (aggregate data shown in main Figure 2B). Shown is the average of two independent repeats. (**D**) Thermal denaturation of mɑQSOX1 and four CUMAb designs using nano differential scanning fluorimetry. Shown is the average of two independent repeats. (**E**) Crystal structure of oxidoreductase fragment of human QSOX1 in complex with parental mouse antibody (QSOX1 in dark gray, parental mouse antibody Fv in light gray, PDB entry 4IJ3) aligned with crystal structure of oxidoreductase fragment of human QSOX1 in complex with hɑQSOX1.4 (QSOX1 in purple, hɑQSOX1.4 Fv in magenta, PDB entry 8AON. Structures are nearly identical (0.7 Å) despite 51 Fv mutations.

**Supplementary Figure 2:**
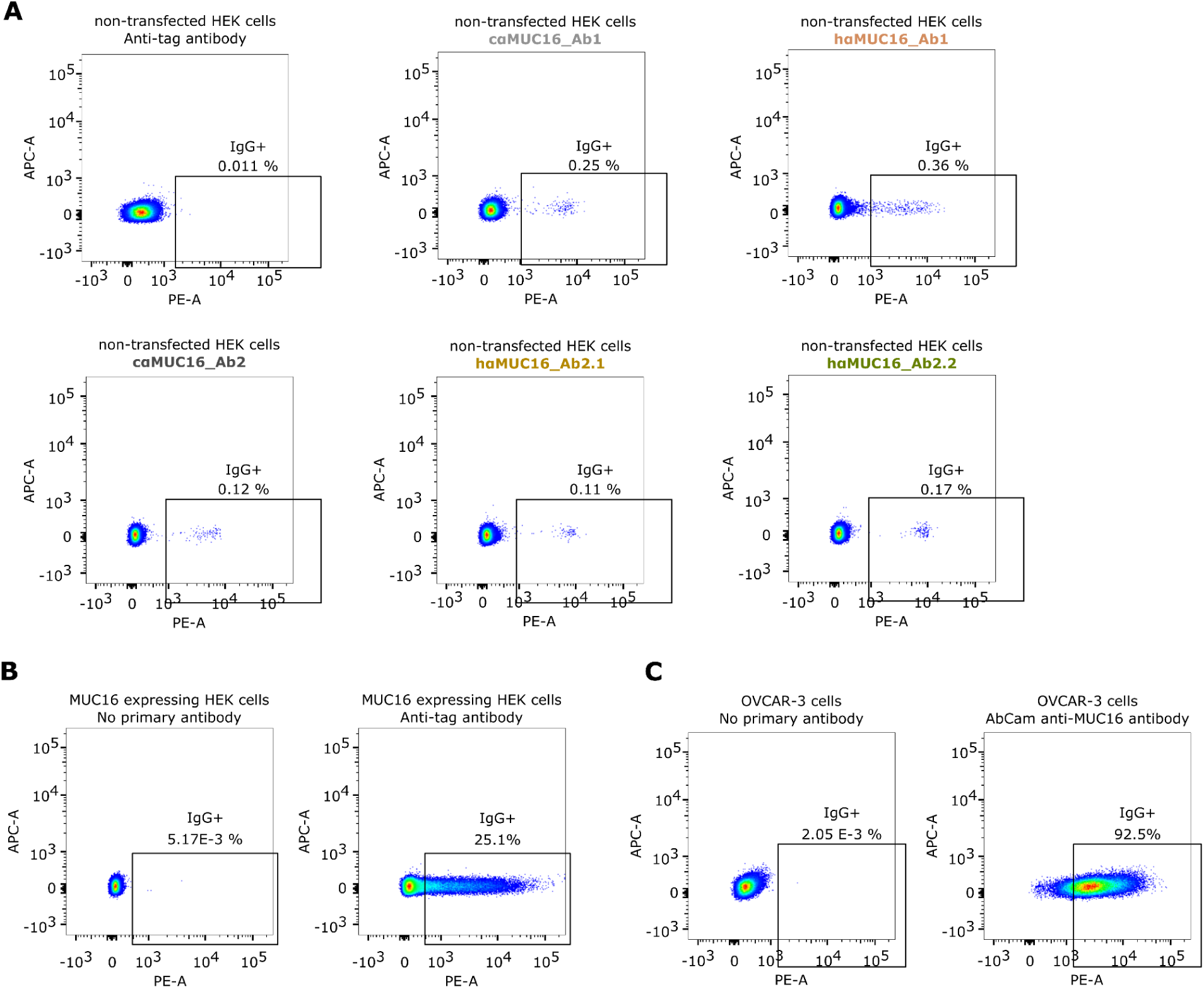
(**A**) Non-transfected HEK293 cells were stained with a commercial anti-tag antibody, cɑMUC16_Ab1, hɑMUC16_Ab1, cɑMUC16_Ab2, hɑMUC16_Ab2.1, and hɑMUC16_Ab2.2, demonstrating that none of the five antibodies showed significant binding to cells not expressing the MUC16 construct. (**B**) HEK293 cells expressing the MUC16 construct were stained either without a primary antibody or with a commercial anti-tag antibody, demonstrating that the secondary antibody alone does not label the cells and that the cells do display the intended construct. (**C**) Acetone-fixed OVCAR-3 cells were stained either without a primary antibody or with a commercial anti-MUC16 antibody, demonstrating that the secondary antibody alone does not label the cells and that the cells do display MUC16.

**Supplementary Figure 3:**
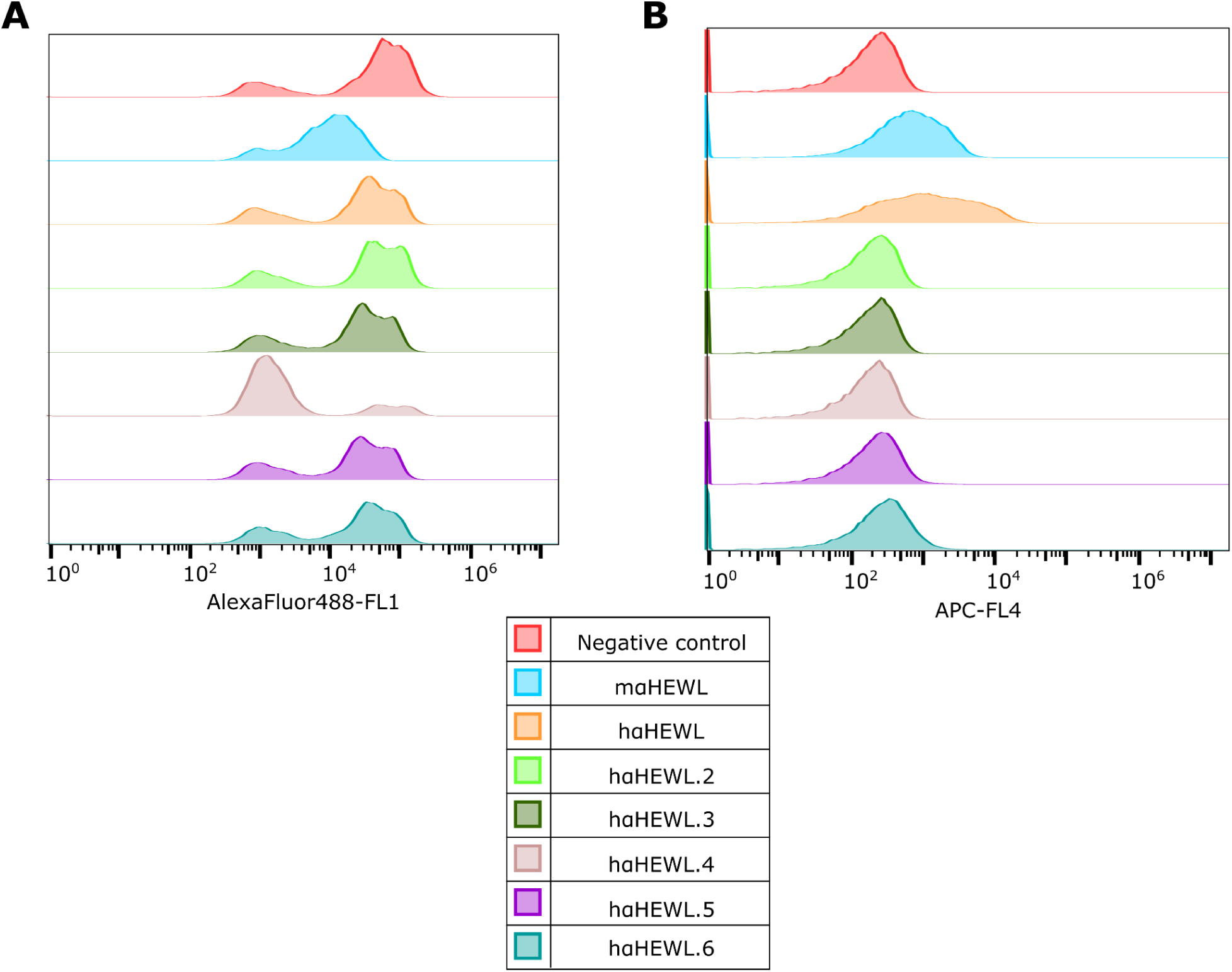
(**A**) Expression levels of negative control (the G6 anti-VEGF^7^ antibody), mouse anti-HEWL antibody (mɑHEWL), and 6 CUMAb designs, all formatted as scFv. (**B**) Binding levels to HEWL of the eight antibodies.

**Supplementary Figure 4:**
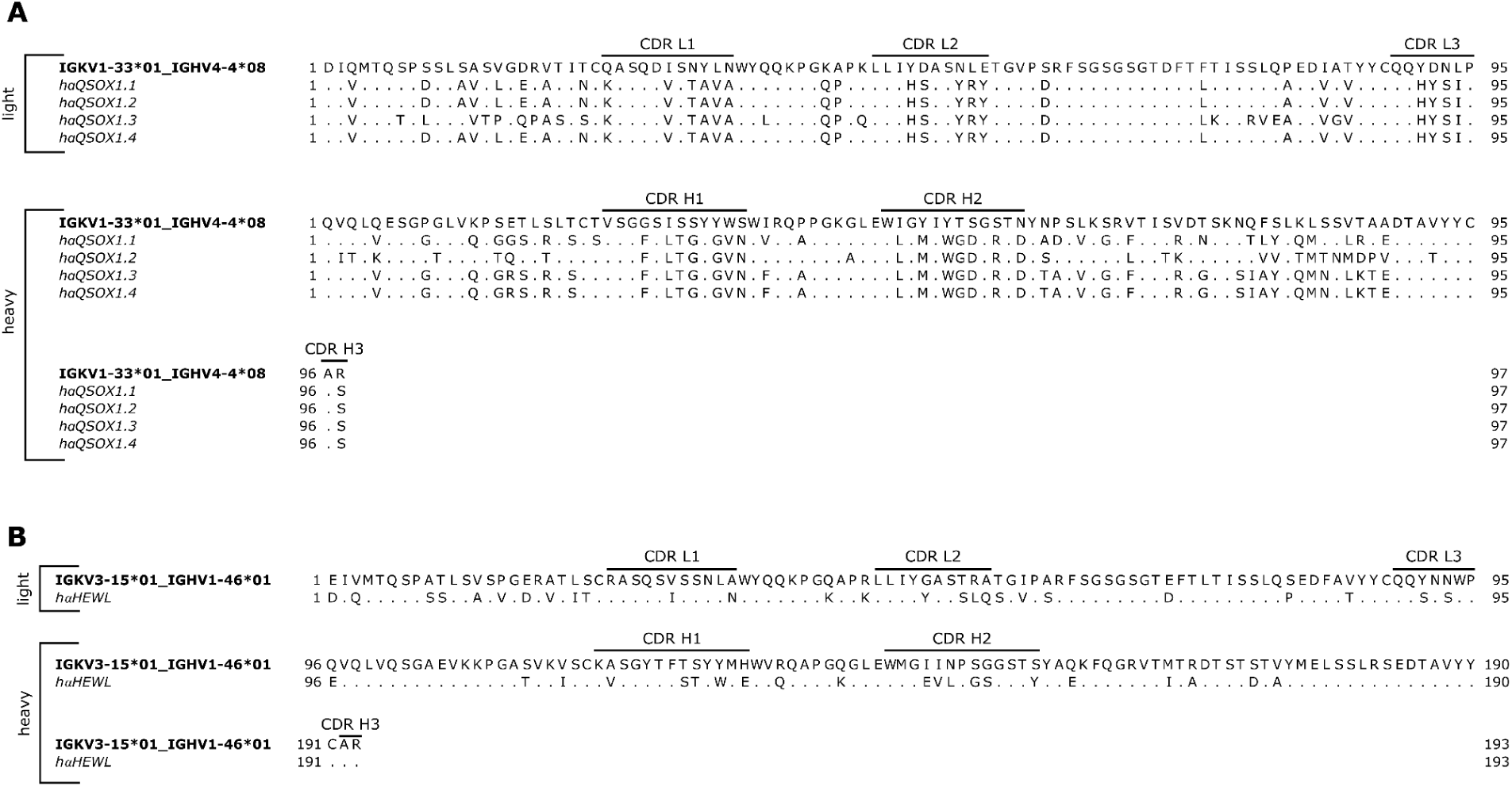
Sequence alignments of V genes in successful (**A**) anti-QSOX1 and (**B**) anti-HEWL designs to the human germline V genes with the highest sequence identity.

**Supplementary Figure 5:**
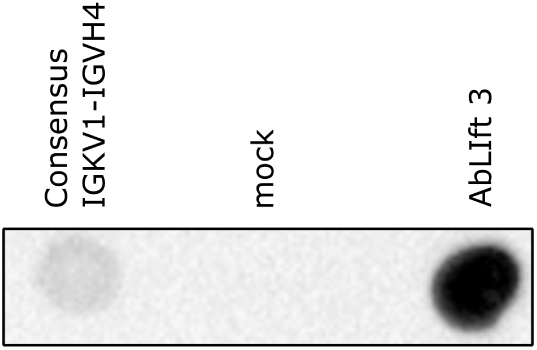
Dot-blot analysis of anti-QSOX1 CDRs grafted onto most homologous consensus frameworks IGKV1-IGHV4 compared to a mock transfection and well-expressed AbLIFT design 3^6^.

